# Inter-individual deep image reconstruction

**DOI:** 10.1101/2021.12.31.474501

**Authors:** Jun Kai Ho, Tomoyasu Horikawa, Kei Majima, Fan Cheng, Yukiyasu Kamitani

## Abstract

The sensory cortex is characterized by general organizational principles such as topography and hierarchy. However, measured brain activity given identical input exhibits substantially different patterns across individuals. Although anatomical and functional alignment methods have been proposed in functional magnetic resonance imaging (fMRI) studies, it remains unclear whether and how hierarchical and fine-grained representations can be converted between individuals while preserving the encoded perceptual content. In this study, we trained a method of functional alignment called neural code converter that predicts a target subject’s brain activity pattern from a source subject given the same stimulus, and analyzed the converted patterns by decoding hierarchical visual features and reconstructing perceived images. The converters were trained on fMRI responses to identical sets of natural images presented to pairs of individuals, using the voxels on the visual cortex that covers from V1 through the ventral object areas without explicit labels of the visual areas. We decoded the converted brain activity patterns into the hierarchical visual features of a deep neural network using decoders pre-trained on the target subject and then reconstructed images via the decoded features. Without explicit information about the visual cortical hierarchy, the converters automatically learned the correspondence between visual areas of the same levels. Deep neural network feature decoding at each layer showed higher decoding accuracies from corresponding levels of visual areas, indicating that hierarchical representations were preserved after conversion. The visual images were reconstructed with recognizable silhouettes of objects even with relatively small numbers of data for converter training. The decoders trained on pooled data from multiple individuals through conversions lead to a slight improvement over those trained on a single individual. These results demonstrate that the hierarchical and fine-grained representation can be converted by functional alignment, while preserving sufficient visual information to enable inter-individual visual image reconstruction.

**Highlights:**

● Neural code converters convert brain activity patterns across individuals with moderate conversion accuracy and learn reasonable visual area correspondences at the same level between individuals.
● The converted brain activity patterns can be decoded into hierarchical DNN features to reconstruct visual images, even though the converter is trained on a small number of data samples.
● The information of hierarchical and fine-scale visual features that enable visual image reconstruction are preserved after functional alignment.

## 1. Introduction

Sensory information is generally thought to be processed through a hierarchical pathway that detects topographically organized simple local features in the early stages and then progressively complex global features in the later stages, leading to holistic perception. In the ventral visual pathway, a stimulus is first processed in the striate cortex (V1) to extract simple features, such as edges (Hubel and Wiesel, 1962), and is then further processed in the extrastriate cortices (V2– V4) and higher visual cortex (HVC) to detect more complex visual features, such as shape and face attributes, eventually identifying objects and scenes (Mishkin and Ungerleider, 1982). Whereas general principles such as topography and hierarchy appear to govern the organization of the visual cortex (VC), individual brains differ substantially in both macroscopic anatomy and the fine-grained organization of feature representations. These individual differences make it challenging to relate visual cortical activity and perceptual content by simple mapping rules common across individuals.

The recent advances in deep neural networks (DNNs) have enabled detailed analyses of hierarchical feature representations across visual cortical areas (Yamins et al., 2014; Güçlü and van Gerven, 2015, 2017; Horikawa and Kamitani, 2017). Previous encoding and decoding studies have shown that DNNs pre-trained on natural images exhibit a correspondence between visual areas and DNN layers. These findings indicate that the visual cortex processes increasingly complex visual features along the ventral neural pathway, similar to how DNNs process image features. Furthermore, perceptual content encoded in brain responses have been successfully reconstructed as images via DNN-based reconstruction algorithms (Shen et al., 2019a and 2019b). The deep image reconstruction (Shen et al., 2019a) first predicts the DNN features of an image from the brain activity given that image as a stimulus, and then an initial image is iteratively optimized such that its DNN features become close to the predicted DNN features. DNN feature decoding enables comprehensive evaluations of hierarchical visual representations, and visual image reconstruction allows holistic evaluations of how accurately perceptual content are encoded in the brain activity patterns. However, these predictive models require training data derived from hours of experiments that measure the brain responses to hundreds or thousands of images. Furthermore, a model trained on one subject does not generalize to other subjects because of individual differences in macroscopic brain structure and fine-grained neural representations.

Methods for the anatomical and functional alignment of different individuals’ brains have been developed in decades of functional magnetic resonance imaging (fMRI) studies to compensate for individual differences. Human brain anatomy differs across individuals in terms of shape, size, and local anatomical landmarks. Functional brain area parcellation that clusters voxels/vertices with similar properties produces similar brain areas on the individual level, but they still exhibit distinct topological features (Blumensath et al., 2013; Laumann et al., 2015). The visual areas delineated by the retinotopy principle (Engel et al., 1994; Sereno et al., 1995) are often similar but not the same across individuals. Anatomical alignment could mitigate the anatomical difference by matching the anatomical features between brains (Fischl et al., 2008; van Essen, 2004, 2005), but it still cannot perfectly align the functional topography across individuals (Watson et al., 1993). Functional alignment adopts an anatomy-free approach by learning statistical relationships between subjects’ brain activity patterns (Haxby et al., 2011; Yamada et al., 2011, 2015; Chen et al., 2015; Bilenko and Gallant, 2016; Guntupalli et al., 2016). Methodologies of functional alignment include pairwise alignments between two subjects, such as a neural code converter (Yamada et al., 2015), and template-based alignments, in which a shared template among subjects is constructed, such as hyperalignment (Haxby et al., 2011). Functional alignment methods have uncovered common neural representations across individuals concealed under substantial individual variations in brain responses. However, those investigations have often focused on a few specific features, such as object categories, image contrast, retinotopy, and semantics (Haxby et al., 2011; Yamada et al., 2015; Bilenko and Gallant, 2016; Van Uden et al., 2018), leaving it unclear whether distinct levels of fine-grained neural representations of hierarchical visual features can be converted across individuals such that an individual’s perceptual experience can be reconstructed using other individuals’ models. Furthermore, the previous studies have separately performed alignments on different brain areas using rough anatomical correspondences across individuals (Güçlü and van Gerven, 2015). It remains unknown whether data-driven methods trained on fMRI data can automatically detect hierarchical representations of distinct levels of visual features common across individuals.

Here, we aim to investigate whether and how fine-grained neural representations of hierarchical visual features can be converted between individuals while preserving the encoded perceptual content. For this purpose, we analyzed the brain activity converted via a functional alignment method (neural code converter; Yamada et al., 2015) using the decoding of hierarchical DNN features (Horikawa and Kamitani, 2017) and reconstruction of perceived images (deep image reconstruction; Shen et al., 2019a). We also adopted other methods of pairwise alignment, including Procrustes transformation (Schönemann, 1966), optimal transport (Bazeille et al., 2019), and a template-based pairwise alignment via hyperalignment (Haxby et al., 2011). Our aim is not to exhaustively evaluate all available methods of pairwise alignment, but to show the robustness of the results across several methods. We restricted the work within the methods of pairwise alignment. We do not discuss the shared template because it is difficult to interpret the correspondence of visual subareas between subjects in a shared template, and the question of how best a template can be estimated is distinct from the alignment methods (Bazeille et al., 2021). We used the template-based alignment only to construct a pairwise transformation via the template, which is called template-based pairwise alignment. We first constructed machine learning-based models that convert an fMRI pattern in the VC of one subject (the source) to the individual voxel responses of another subject (the target) given identical sequences of natural image stimuli (Fig. 1A). We also train DNN feature decoders with measured fMRI responses of the target subject (Fig. 1A). Then, given the source subject’s brain responses to novel stimuli, the converter transforms the brain activity into the target brain space (Fig. 1B). The converted brain activity is decoded by the DNN feature decoders pre-trained on the target subject, and then the decoded features are used in a reconstruction algorithm to create images (Fig. 1B).

**Fig. 1.**
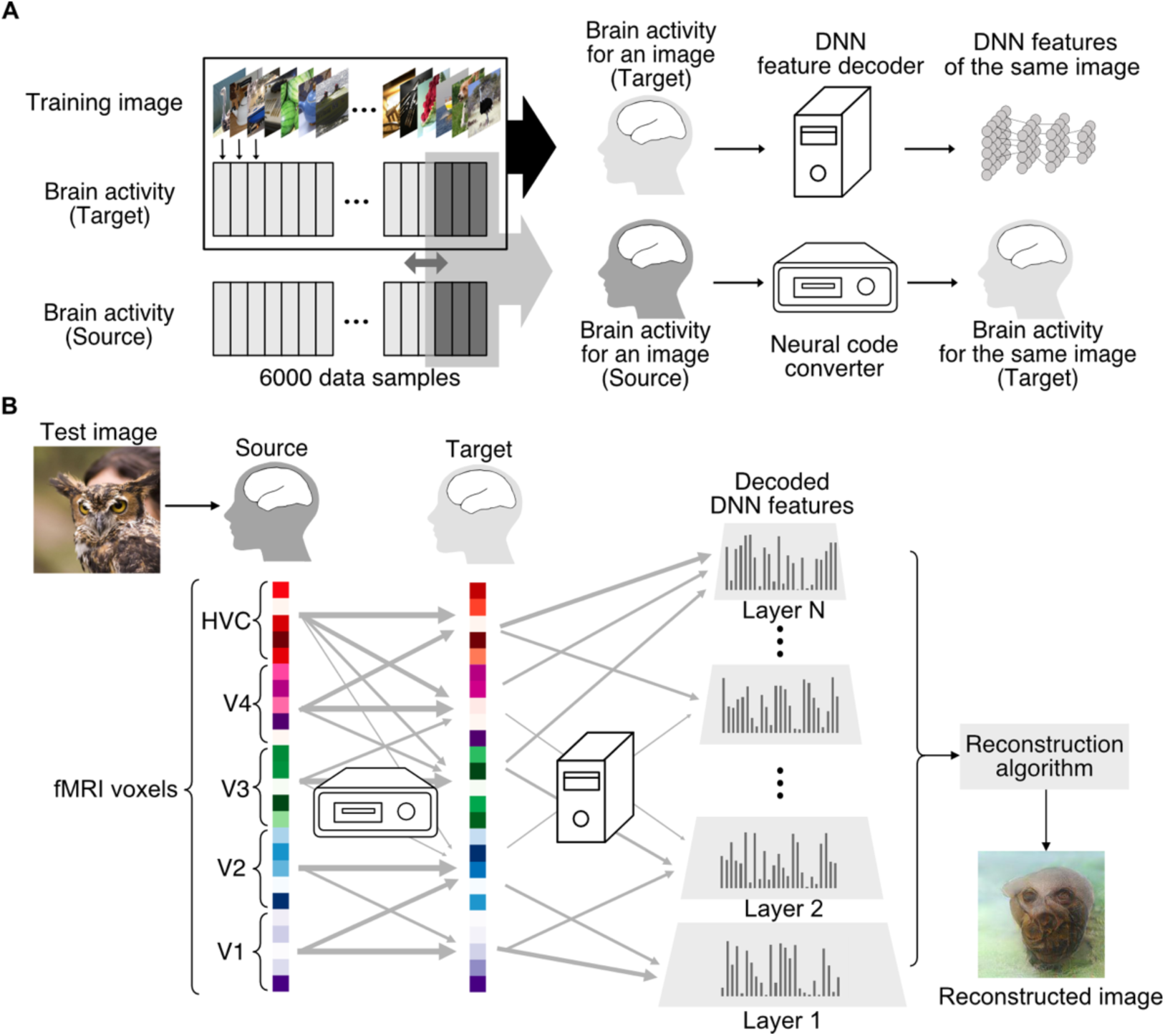
Inter-individual deep image reconstruction. (A) Training of the DNN feature decoders and a neural code converter. DNN feature decoding models were trained on the 6,000 samples of measured fMRI activities of the target subject and the corresponding DNN features. A converter model is trained on a subset of 6,000 samples of fMRI data responses to an identical stimulus sequence from both the source and target subject. No explicit information about cortical hierarchy is provided at the training stage. (B) Inter-individual DNN feature decoding and visual image reconstruction. The converter model converts the source subject’s stimulus-induced fMRI pattern to the target subject’s brain space. The converted fMRI pattern is then decoded (or translated) into DNN feature pattern using the feature decoders. Finally, the decoded features are fed into the reconstruction algorithm to reconstruct the stimulus image perceived by the source subject.

In this study, we first show that machine learning-based converter models automatically learn the hierarchical correspondence of visual subareas between subjects even though explicit information about cortical hierarchy is not given in the training. DNN feature decoding from the converted fMRI responses at each hierarchical DNN level shows greater accuracy in the corresponding levels of visual subareas than in other levels, indicating that fine-grained hierarchical feature representations are preserved. Visual image reconstruction using the decoded DNN features from the converted fMRI responses produces faithful reconstructions of the viewed images even with small numbers of data for the converter training. We also demonstrate that the information about cortical hierarchy used in the training does not improve the performance of converters, given sufficient training data. Lastly, we pool data from multiple subjects by neural code conversions and show that DNN feature decoders trained on the pooled data achieve a slight improvement in the inter-individual visual image reconstruction. These results demonstrate that the hierarchical correspondence can be automatically detected and the fine-grained representations of visual features can be preserved across individuals by the neural code converters, providing an efficient way to create visual image reconstructions even for novel individuals.

## 2. Results

### 2.1 fMRI data

We analyzed fMRI data of the five subjects in the previously published studies (Shen et al., 2019a; Horikawa and Kamitani, 2022). For two of the five subjects, we additionally collected data for the present study (see Materials and Methods: “fMRI datasets”). The dataset consisted of fMRI data measured when subjects viewed images, each presented in an 8-s block (four fMRI volumes). To acquire training data, the presentation of 1,200 natural images was repeated five times. For test data, the presentation of 50 natural images was repeated 24 times in the test natural image session, and the presentation of 40 artificial images was repeated 20 times in the test artificial image session. Artificial images (simple geometric shapes) were introduced to examine how models trained on natural images generalize to a different type of images. The fMRI data were averaged in each 8-s stimulus block (four fMRI volumes shifted by 4 s to account for hemodynamic delays). Thus, 6,000 (5 × 1,200) training samples, 1,200 (24 × 50) test samples with natural images, and 800 (20 × 40) test samples with artificial images were available. In decoding and reconstruction analyses, test samples were further averaged across repetitions (blocks) for each image. Note that the fMRI data collection for 6,000 training samples required approximately 800 minutes of scan sessions in each subject, which were performed on several different days. Although some of the training data and the test data were collected at different times separated by more than several months or even a year, the trained model generalized well across the datasets as demonstrated in Shen et al. (2019a).

### 2.2 Neural code conversion

We first examined how the results of neural code conversion reflect cortical hierarchy using several evaluation methods. We constructed a neural code converter model between each pair of subjects, using one as the target subject and the other as the source subject, which resulted in 20 individual pairs. A converter model comprises a set of regularized linear regression models (ridge regression), each trained to predict the activity of each voxel of the target subject’s brain from the source subject’s brain activity pattern in a broad region of interest (ROI) that covered the lower to higher visual cortex termed VC (see Materials and Methods: “Methods of functional alignment”). In the current study, neural code converter models were trained using a varying number of training samples (300, 600, 900, 1,200, 2,400, 3,600, 4,800, or 6,000 samples). Unless otherwise noted, we show the results obtained using 2,400 training samples (two repetitions of 1,200 images) as a representative case.

VC consists of V1–V4 and ventral object-responsive areas (see Materials and Methods: “Regions of interest”). We defined the continuous region covering the lateral occipital complex (LOC), fusiform face area (FFA), and parahippocampal place area (PPA) as the higher visual cortex (HVC). In the analyses of this section, all VC voxels were used as inputs to the converter without additional voxel selection (see Materials and Methods: “Neural code converter”). Conversion results were evaluated in individual ROIs (subareas) in the target subject’s brain space.

Although we mainly present the results from the neural code converter analysis (Yamada et al., 2015), we also performed similar pairwise alignment analyses using Procrustes transformation (Schönemann, 1966), optimal transport (Bazeille et al, 2019), and template-based pairwise alignment via hyperalignment (Fig. S1; see Materials and Methods: “Methods of functional alignment”) to confirm the robustness of the results across different functional alignment methods (for an evaluation of different methods, see Bazeille et al. [2021]). Compared to other methods, the neural code converter is simple and less computationally expensive.

We evaluated the models using two methods: (a) pattern correlation, which is the spatial Pearson correlation coefficient between the converted and measured voxel patterns for a test image, and (b) profile correlation, which is the Pearson correlation coefficient between the sequences of converted and measured individual voxel responses to the 50 natural test images (Fig. 2A). The pattern correlation for an image was defined by the mean of 24 samples (converted) × 24 samples (measured) = 576 correlation coefficients. The profile correlation for each voxel was defined by the mean of 24 repetitions (converted) × 24 repetitions (measured) = 576 correlation coefficients. The obtained correlation coefficients were normalized by their noise ceilings to account for the noise in fMRI brain responses over repeated measurements with the same stimulus (Hsu et al., 2004; Lescroart and Gallant, 2019; see Materials and Methods: “Noise ceiling estimation”). To summarize the results, the correlation coefficients were further averaged across images and voxels for the pattern and the profile correlations, respectively, in each individual pair and each ROI.

**Fig. 2.**
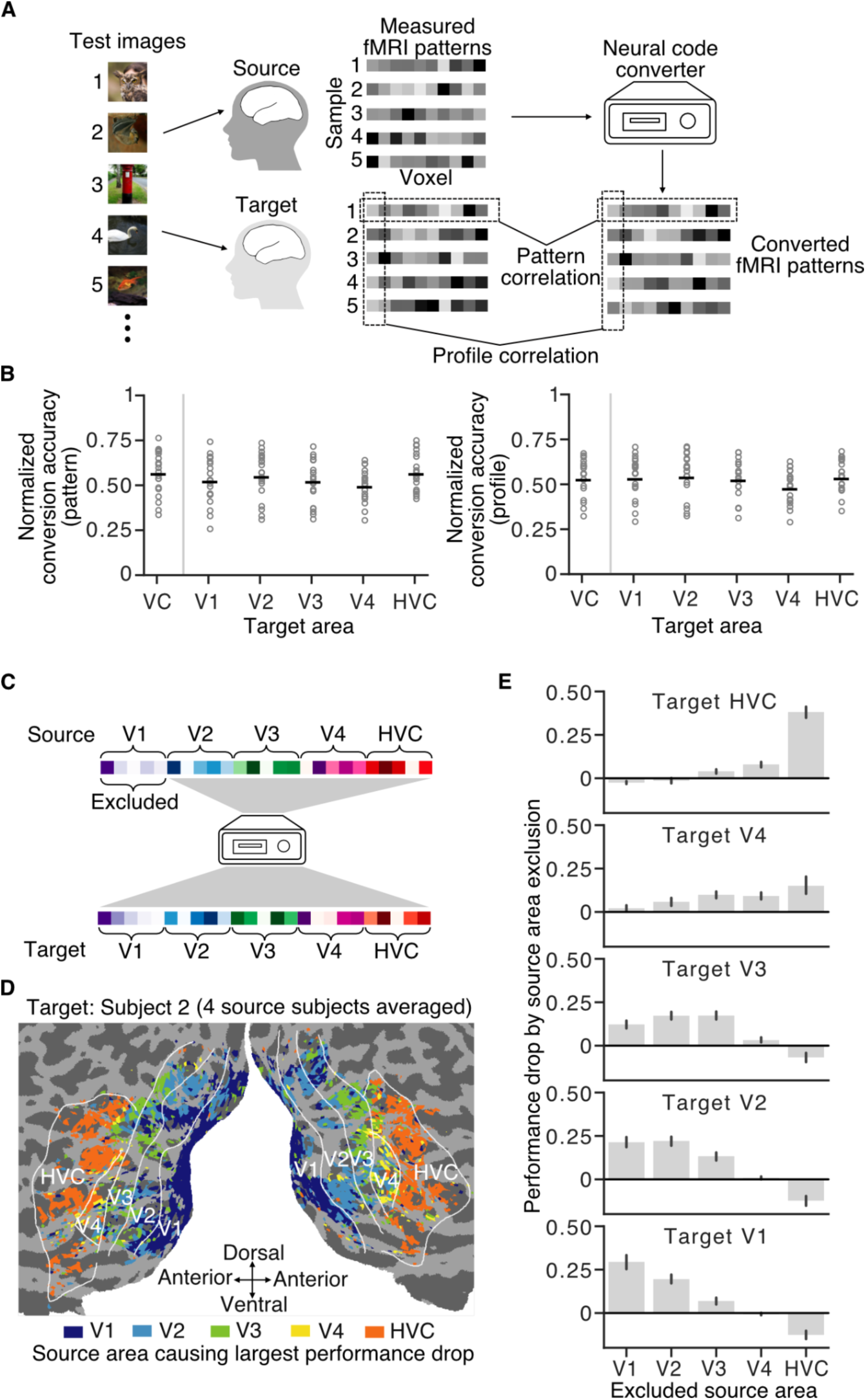
Performance of neural code converters and cortical hierarchical correspondence. (A) Evaluations of neural code converters. Two evaluations were performed by computing the Pearson correlation coefficients: pattern and profile correlations. (B) Conversion accuracy. Distributions of the normalized pattern or profile correlation coefficients of 20 individual pairs are shown for the VC and visual subareas. Each horizontal black dash indicates the mean value; each circle represents the correlation coefficients of an individual pair. (C) Ablation analysis on neural code converters. The analysis was performed by excluding one source visual area from the prediction of target voxel activities. (D) Cortical map of the effects of source area exclusion. The cortical map is shown for one target subject (Subject 2). Each voxel on the target brain is colored by the index of the excluded visual area that caused the largest performance drop when testing with the natural image test dataset (performance drops were averaged across four source subjects for a single target subject; see Fig. S3 for other target subjects). Only voxels that generate reliable responses with noise ceilings above a threshold are shown (see Materials and Methods: “Noise ceiling estimation”). (E) Mean performance drop caused by source area exclusion. Each bar represents the mean performance drop averaged across voxels in a target area when a source area was excluded during prediction (averaged over 20 individual pairs; error bars, 95% confidence interval [C.I.] from 20 individual pairs).

While we primarily performed analyses on the samples within a conversion pair (Smith et al. 2018), group results, where each data point represents an individual pair, are shown in main figures for illustrative summary purposes. The normalized pattern correlation coefficients in individual pairs are shown for different ROIs of the target subject in Fig. S2A (left), and their distributions across all conversion pairs are shown in Fig. 2B (left). The mean normalized pattern correlation for the whole VC was 0.56 ± 0.06 (mean with 95% C.I.) over 20 individual pairs, with the visual subareas showing comparable distributions. Examples of converted brain activity patterns are shown with the targets’ brain activity patterns in Fig. S2B. The mean normalized profile correlation for VC was 0.53 ± 0.05 over 20 individual pairs (Fig. S2A right for individual pairs; Fig. 2B right for group results). The subareas also yielded distributions similar to those of the VC. The conversion accuracy was modest in both pattern and profile correlations across all visual subareas but comparable to those in the previous study (Yamada et al., 2015). Other methods of functional alignment showed similar conversion accuracies, with optimal transport showing higher accuracies (Fig. S3).

To see how the source visual areas contributed to the conversion accuracy for each voxel in each target visual area, we excluded one of the source visual subareas (V1, V2, V3, V4, or HVC) from the inputs to the trained converter model (Fig. 2C). We evaluated the drop in performance (normalized profile correlation difference) relative to the performance of the condition using all source visual subareas (*i.e.*, the whole VC). This ablation analysis revealed that the effects of the source area exclusions varied with the area in the target brain. The largest drop in performance of a target voxel was often caused by the exclusion of the corresponding source area (Fig. 2D, target S2; see Fig. S4A for the other subjects). On average, the peak of performance drop shifted from lower to higher excluded source areas along the hierarchy of the target areas (Fig. S4B for results of some individual pairs; Fig. 2E for group results). The results indicate that the machine learning-based neural code converter models automatically detect a “low-to-high” hierarchical correspondence between source and target visual areas even without explicit information about the anatomy.

### 2.3 DNN feature decoding

We next used DNN feature decoding analysis (Horikawa and Kamitani, 2017) to examine whether fine-grained representations of visual features were preserved in the converted fMRI activity patterns. Feature decoders had been trained to predict the DNN feature values of the stimuli using 6,000 training samples of a target subject’s fMRI activity patterns in the whole VC and individual visual subareas. The feature decoders were applied to the converted brain activities to predict the DNN features of the test images (“Across-functional” condition; see Materials and Methods: “DNN feature decoding analysis”). Following the original paper (Shen et al., 2019a), we used the average fMRI data over the repetitions for each test image as the input to feature decoders. The decoding accuracy of each DNN unit was obtained by calculating the Pearson correlation coefficient between the sequences of the decoded and true feature values for the test images. We further took the mean decoding accuracy over all DNN units in each layer.

For comparison, we performed the same analysis with anatomically aligned brain activity. The source subject’s fMRI images were aligned to the target’s anatomical template and then used for DNN feature decoding (“Across-anatomical”; see Materials and Methods: “Anatomical alignment”). We also compared the results with the standard within-individual decoding, in which DNN features were predicted using the decoders trained on the same subject’s data (“Within”).

We first evaluated feature decoding performance obtained from the whole VC (in the target space) of the converted fMRI activity. The results of the neural code converter (Across-functional) showed a lower but comparable performance with the within-individual results, with similar trends across layers, both in individual pairs and at the group level (Fig. S5A for results of individual pairs; Fig. 3A for group results). Anatomical alignment (Across-anatomical) performed the worst among the three conditions, with accuracies below 0.1 in most layers, both in individual pairs and at the group level. The results show that the neural code converters have an advantage over anatomical alignment in DNN feature decoding. Other methods of functional alignment showed similar DNN decoding accuracies, with optimal transport showing lower accuracies (Fig. S6).

**Fig. 3.**
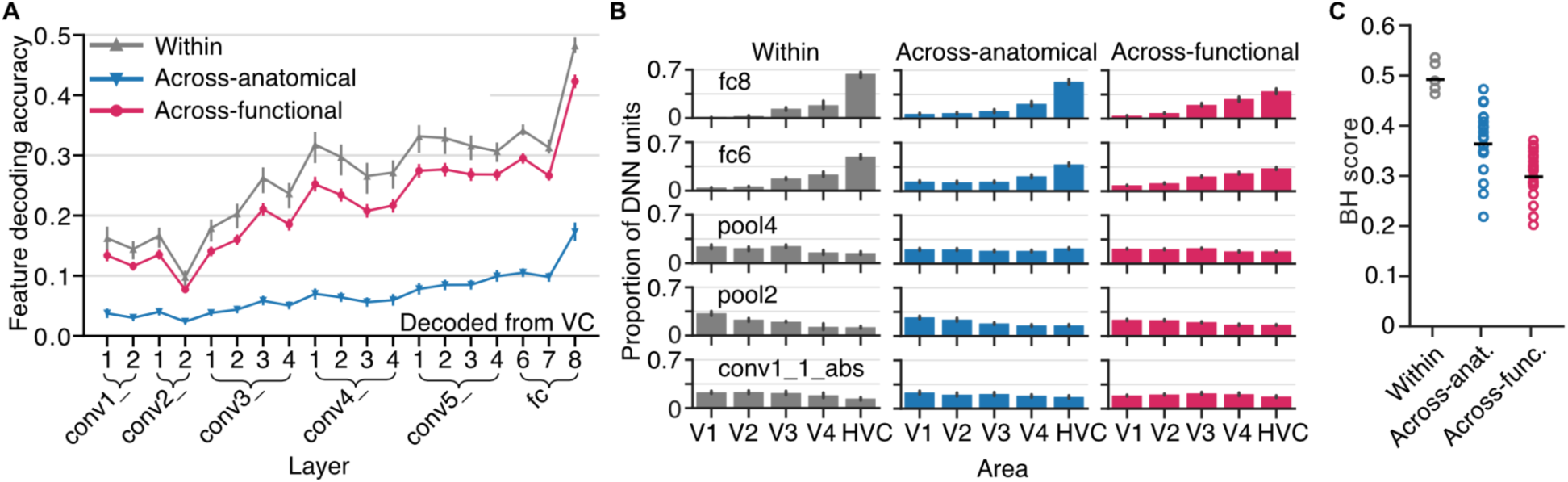
DNN feature decoding and hierarchical representation. (A) DNN feature decoding accuracy from the whole visual cortex (VC). Decoding accuracies for each layer of the VGG19 model are shown for the Within, Across-anatomical, and Across- functional conditions (error bars, 95% C.I. from five subjects for the Within condition, and from 20 individual pairs for the Across-anatomical and Across-functional conditions). (B) Proportion the “top visual area” (best decodable area for each DNN unit) across DNN units in each layer. Only five representative layers are shown. Each bar indicates the mean proportion of DNN units over five subjects for the Within condition or over 20 individual pairs for the Across-anatomical and Across-functional conditions (error bars, 95% C.I. from five subjects or 20 pairs.). (C) Brain hierarchy (BH) score. The horizontal black dashes indicate the mean BH score over subjects or pairs; each circle represents the BH score for a subject or a pair. though the converter was blind to cortical hierarchy information during the training.

We next performed decoding analysis on each DNN unit using voxels from individual visual areas (V1–V4 and HVC in the target space) and identified the visual area that gave the highest decoding accuracy for each unit (“top visual area”), as in Nonaka et al. (2021). We then computed the distribution of the top visual area across DNN units in a given layer. We observed a shift of the peak area, from lower to higher areas, along the DNN hierarchy in all conditions (Fig. S5B for results of individual pairs; Fig. 3B for group results). To quantify the degree of hierarchical correspondence between brain areas and DNN layers, we used the decoding-based brain hierarchy (BH) score (Nonaka et al., 2021), which is based on the rank correlation between the hierarchical levels of the DNN layer and the top brain area across DNN units (Fig. 3C; see Materials and Methods: “Brain hierarchy (BH) score”). The results of the within-individual condition replicated the previous findings with a BH score of around 0.5 (Horikawa and Kamitani, 2017; Nonaka et al., 2021). Although anatomical alignment (Across-anatomical) showed overall low accuracies in feature decoding (Fig. 3A), the hierarchical correspondence is largely preserved when quantified by the BH score (Fig. 3B, C). This is presumably because anatomical alignment maps a macroscopic organization of hierarchical visual areas between subjects and the relative amount of information is preserved. The inter-individual conversion (Across-functional) showed a lower but substantial degree of hierarchical correspondence even

### 2.4 Visual image reconstruction

After confirming that multiple levels of DNN feature representations can be decoded from converted brain activity, we next sought to determine if we could reconstruct visual images via DNN features decoded from converted brain activity (deep image reconstruction, Shen et al., 2019a; see Materials and Methods: “Visual image reconstruction”). In addition to the natural images, we also performed the reconstruction analysis on the artificial images of simple geometric shapes (see Materials and Methods: “fMRI datasets”).

We first show examples of the reconstructions from VC for the Within, Across-anatomical, and Across-functional conditions (Fig. 4A). The reconstructed images obtained in the Within and Across-functional conditions captured the characteristics of the presented images, including the shapes and colors of the objects, while reconstructions with anatomical alignment (Across-anatomical) showed neither a recognizable shape nor color of the objects in the presented images (see Fig. S7 and Fig. S8 for other examples of natural images and artificial images). Other methods of functional alignment also produced similar reconstructions, but optimal transport slightly underperformed compared with others (Fig. S9). Here, we only present reconstructions from the averaged fMRI data over all the repetitions (24 and 20 repetitions for natural and artificial images, respectively). The results with the average of different numbers of repetitions are available in the supplemental information (Fig. S10). Even fMRI data of only one repetition could produce discernible reconstructions, with the visual quality increasing with the repetition.

**Fig. 4.**
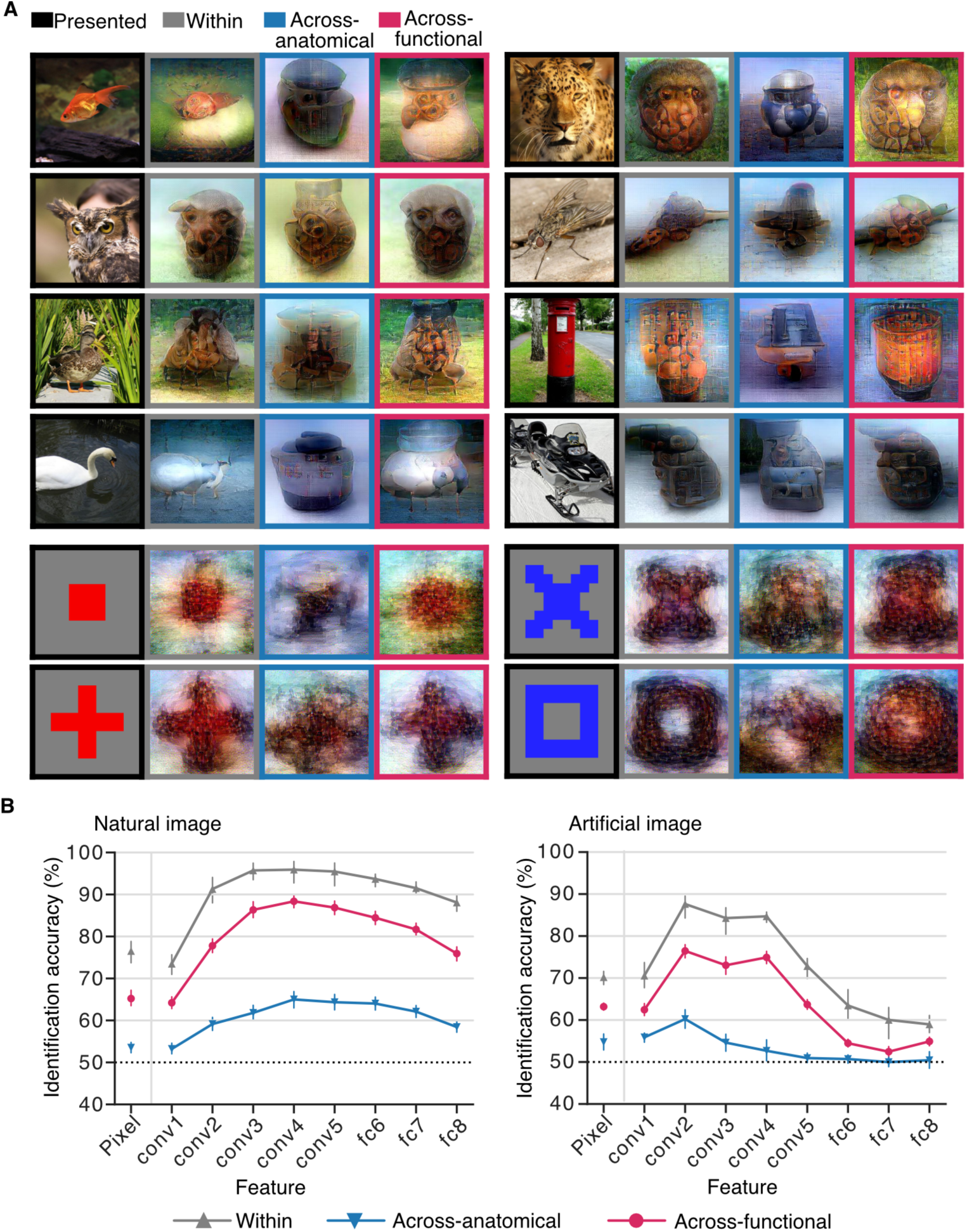
Reconstructed images and evaluations. (A) Within and across-individual reconstructions from the whole visual cortex (VC). The reconstructions under the three analytical conditions for each stimulus image were all from the same source subject. The results for different stimulus images are from different source subjects. (B) Identification accuracy based on pixel values and extracted DNN feature values. A mean identification accuracy was calculated over all reconstructed images for each subject or individual pair. DNN features of images were extracted from the eight layers of the AlexNet model (left, natural images; right, artificial images; error bars, 95% C.I. from five subjects or 20 pairs; dotted lines, chance level = 50%).

For quantitative evaluation, we performed a pairwise identification analysis in which the pixel or DNN feature pattern of a reconstruction was used to identify the true stimulus between two alternatives by choosing the one with a more correlated pattern (see Materials and Methods: “Identification analysis”). DNN feature patterns were extracted using the AlexNet model (Krizhevsky et al., 2012), which is different from the DNN used in our reconstruction method (VGG19 model). The identification was repeated for multiple false alternatives to obtain the accuracy for each reconstruction. For group analysis, the mean identification accuracy was calculated over all reconstructions in each pair. While the within-individual condition (Within) showed overall superior performance both for natural and artificial images, neural code conversion (Across-functional) greatly outperformed anatomical alignment (Across-anatomical) both in individual pairs and at the group level (Fig. S11 for individual pairs; Fig. 4B for group results).

### 2.5 Visual subarea-wise conversion

To examine whether explicit information about cortical hierarchy given in the converter training could improve visual image reconstruction, we performed subarea-wise conversion that predicted the activity values of a voxel in a target area only from the source subject’s corresponding source area (Fig. 5A). All individual pairs showed comparable conversion accuracies to the whole VC conversion, with the mean pattern correlation being 0.58 ± 0.07 and the mean profile correlation being 0.55 ± 0.06 for VC (Fig. S12A for individual paris; Fig. S12B for group results). We then compared the subarea-wise and the whole VC conversions with DNN feature decoding and visual image reconstruction (*c.f.*, Fig. 3 and 4).

**Fig. 5.**
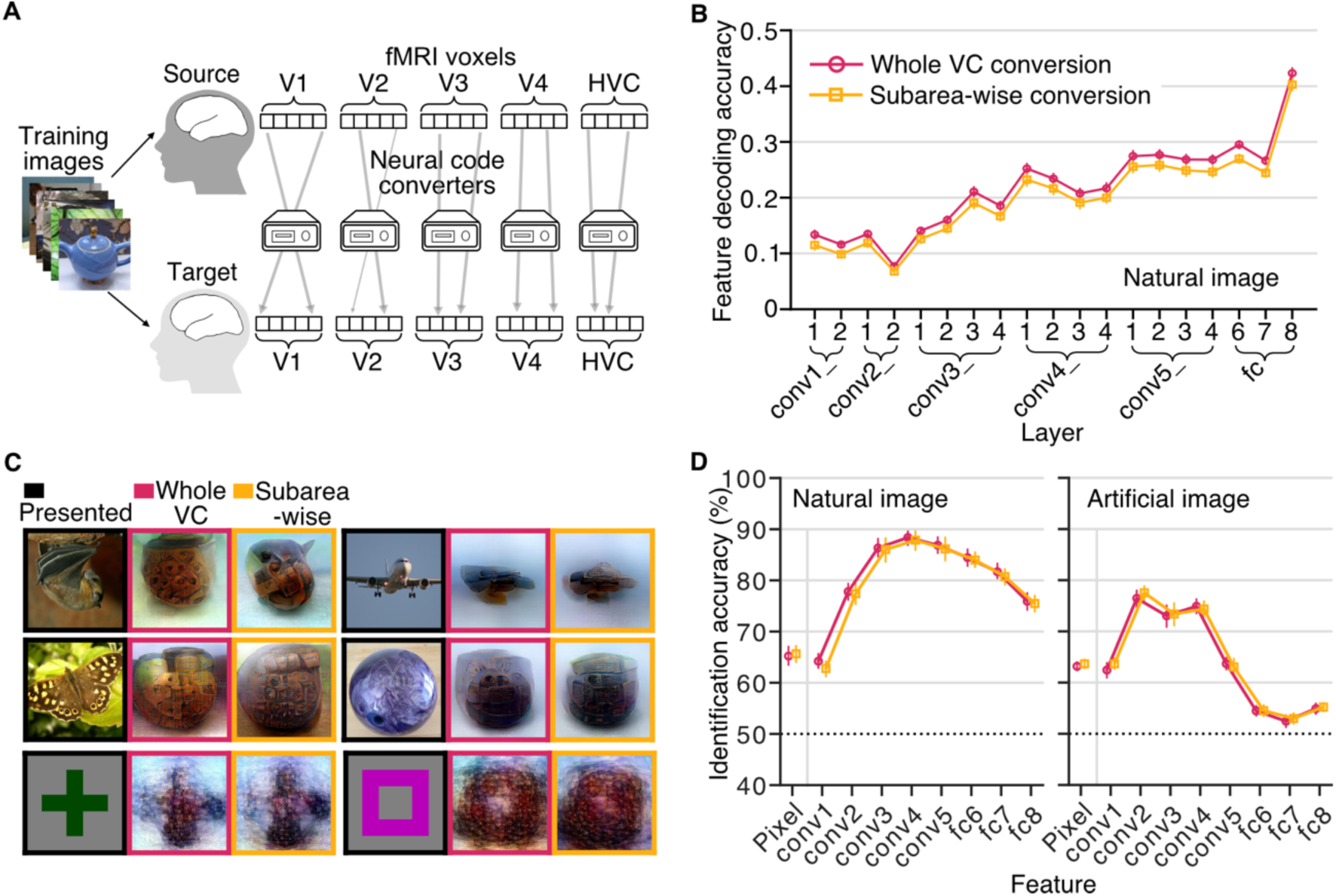
Whole VC vs. subarea-wise conversion. (A) Illustration of the subarea-wise conversion. A converter model was trained on a set of fMRI responses to an identical stimulus sequence. Activity values of a voxel in a target area were predicted only from source subject brain activity patterns in the voxel’s corresponding source area. (B) DNN feature decoding accuracy. The whole VC conversion and subarea-wise conversion were evaluated using DNN feature decoding of natural images (error bars, 95% C.I. from 20 individual pairs). (C) Reconstructed natural and artificial images. (D) Identification accuracies based on pixel values and extracted DNN feature values. DNN features of images were extracted from the eight layers of the AlexNet model (left, natural images; right, artificial images; error bars, 95% C.I. from 20 individual pairs; dotted lines, chance level = 50%).

In the DNN feature decoding of the natural images, the subarea-wise conversion showed similar but slightly lower decoding accuracy than the whole VC conversion across layers in all individual pairs (Fig. S12C) and at the group level (Fig. 5B; ANOVA on the means of individual pairs, effect of conversion type with the DNN layer as a between-subject factor, *F*(1, 361) = 1959, *p* < .001, 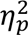 = .84; see Materials and Methods: “Statistics”). Similar results were obtained with the artificial images in some individual pairs and at the group level (Fig. S12C for individual pairs; Fig. S12D for group results; ANOVA on the means of individual pairs, *F*(1, 361) = 260.6, *p* < .001, 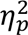 = .42).

Reconstructed images obtained from subarea-wise conversions had a visual quality similar to those of the whole VC conversions (Fig. 5C). In the identification analysis of the natural images (Fig. 5D), only 2/20 pairs showed significantly higher accuracies for the subarea-wise conversion; 6/20 pairs showed higher significant accuracies for the whole conversion (Fig. S12E; ANOVA in individual pairs; effect of conversion type with the DNN layer feature as a between-subject factor). At the group level, the subarea-wise conversion showed lower accuracies (Fig. 5D; ANOVA on the means of individual pairs, *F*(1, 171) = 11.2, *p* < .001, 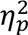 = 0.062). In the identification analysis of the artificial images, 3/20 pairs showed higher significant accuracies for the subarea-wise conversion; 2/20 pairs showed higher significant accuracies for the whole conversion (Fig. S12E; ANOVA in individual pairs), while no statistical difference was found at the group level (Fig. 5D; ANOVA on the means of individual pairs, *F*(1, 171) = 3.87, *p* =.051, 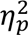= .022). These results indicate that explicit information about the cortical hierarchy does not seem to contribute to the improvement of visual image reconstruction. Rather, the flexibility of the mapping with the whole VC conversion could be beneficial as indicated by the slightly superior performance with the natural images.

### 2.6 Varying the number of training data

One of the potential benefits of the inter-individual analysis is to reduce the number of data required for model training from novel test (source) subjects by using training data from other individuals. The results of the inter-individual analysis so far were obtained using 2,400 samples for converter training, we here investigated how the number of training samples affects the image reconstruction quality by varying the number of data used for the converter training (300, 600, 900, 1,200, 2,400, 3,600, 4,800, and 6,000 training samples) while using all data of the target subject for decoder training (6,000 samples). We also compare the results between the whole VC and the subarea-wise conversions.

While the visual quality of reconstructions decreased with fewer training samples, the reconstructions obtained via converters trained on 300 samples still produced discernible images both in the whole VC and the subarea-wise conversions (Fig. 6A; similar results were obtained for the artificial images, see Fig. S13A). This result indicates that image reconstruction using converters with a small number of training data is feasible, without the need for collecting a full set of fMRI data for a subject.

**Fig. 6.**
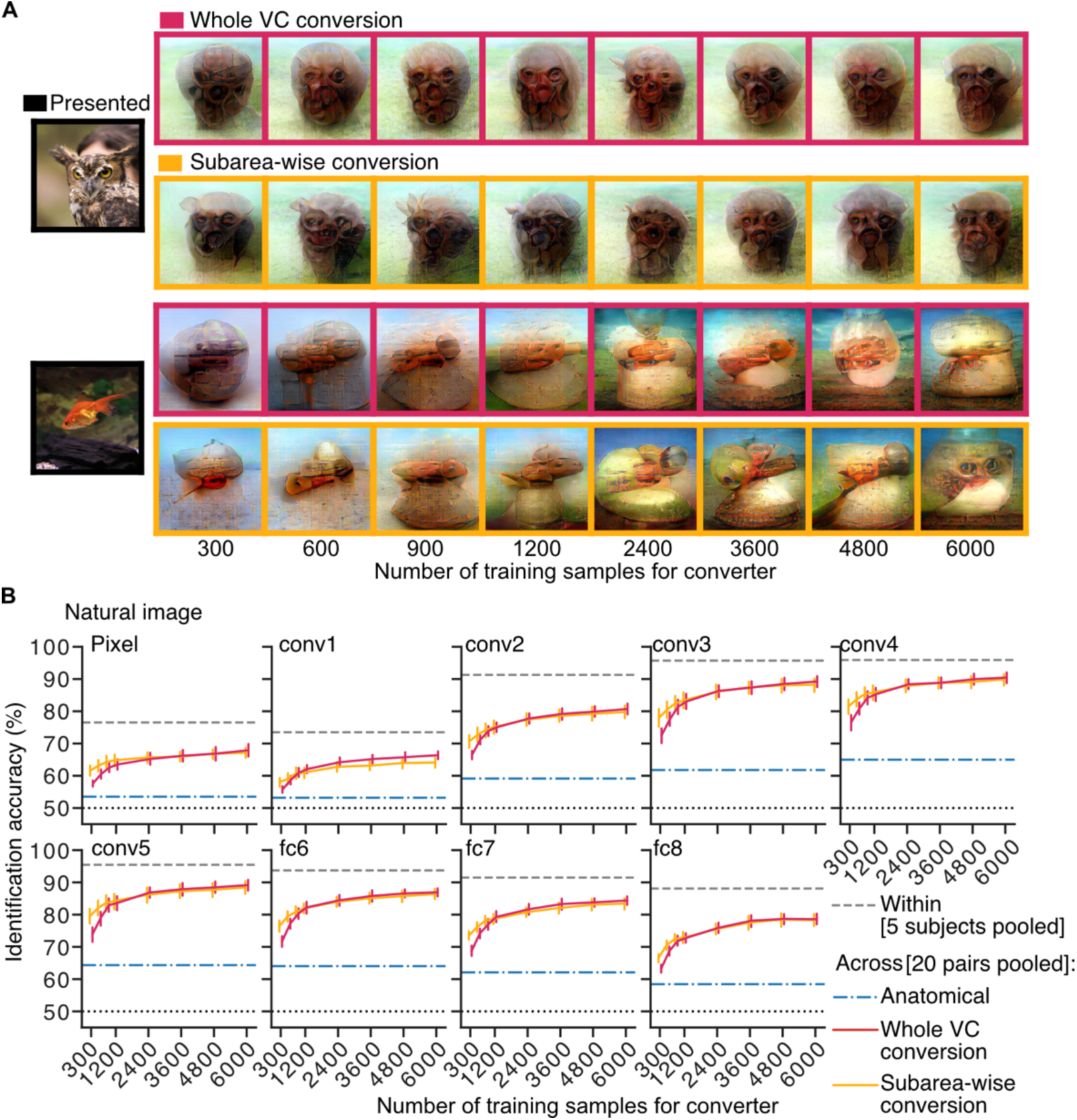
Effect of the number of training data for the converter. (A) Reconstructed images. All reconstructed images were produced from the same subject pair (source: Subject 2, target: Subject 3) (B) Identification accuracy. Identification accuracies were calculated with the pixel values and the extracted DNN feature values (AlexNet) from the reconstructed natural images with varying numbers of training data for the whole VC and subarea-wise converters. The results are shown with those from the within-individual condition (Within) and the anatomical alignment (Across- anatomical) (error bars, 95% C.I. from 20 individual pairs for whole VC and subarea-wise conversions; dotted lines, chance level = 50%).

The identification accuracies increased with the number of training samples, approaching the accuracy of the within-individual (Within) condition (see Fig. S14A for individual pairs and Fig. 6B for group results). The subarea-wise and whole VC conversions showed similar accuracies with more than 1,200 training samples, but the subarea-wise conversion outperformed the whole VC conversion with 1,200 or fewer training samples (ANOVA within individual pairs at each training sample number, effect of conversion type, *p* < .05 in 18, 10, 4, 3, 2, 1, 4, and 1 out of 20 pairs for the eight training sample numbers, respectively; group analysis on the mean accuracies of individual pairs, *p* < .05 at 300, 600, and 900 samples; Bonferroni-corrected by eight). Similar results were obtained for the artificial images (Fig. S14B for individual pairs; ANOVA within individual pairs, effect of conversion type, *p* < .05 in 10, 8, 5, 2, 2, 1, 1, and 2 out of 20 pairs for the eight training sample numbers, respectively; Fig. S13B for group results; group analysis on the mean accuracies of individual pairs, *p* < .05 at 300, 600, and 900 samples; Bonferroni-corrected by eight). Overall, the explicit information about cortical hierarchy does not generally improve reconstruction, but it is beneficial when the number of training data is limited.

### 2.7 Pooling data from multiple subjects

Finally, because the neural code conversion allowed us to pool the data of multiple subjects into a single target subject brain space, we examined the pooling effect on the performance of the inter-individual visual image reconstruction. For a pair of a source and a target subject, we pooled all data from the other three subjects into the target brain space (4 subjects ✕ 6,000 samples = 24,000 samples in total; whole VC conversion; Fig. 7A). We re-trained the decoders on such pooled data and called the decoders “multiple-subject feature decoders,” in contrast to the “single-subject feature decoders,” which were trained on the target subject in the native brain space. For the training of the neural code converter between the source subject’s data and the pooled data, 2,400 samples of the source subject were paired with each set of 2,400 samples from the four pooled subjects. The converted brain activity from the source subject underwent DNN feature decoding with the multiple-subject feature decoders and then visual image reconstruction. The results were compared with those from the single-subject feature decoders.

**Fig. 7.**
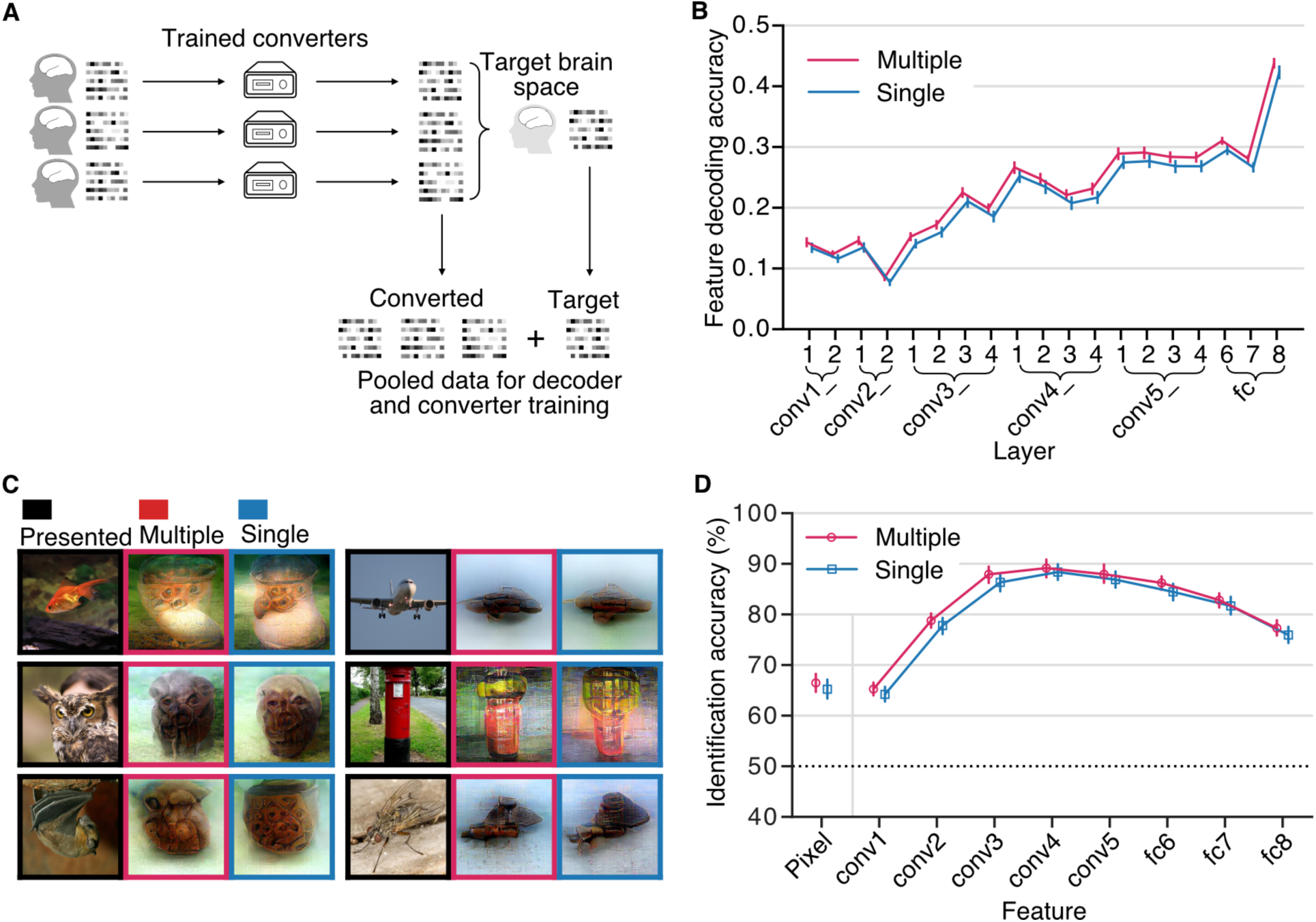
Pooling data from multiple subjects. (A) Illustration of the pooling procedure. For a pair of a source (not shown) and a target subject, the training data of the other three subjects were converted into the target subject’s brain space and DNN feature decoders were re-trained on the converted data of the three subjects plus the target subject’s data (24,000 samples). (B) DNN feature decoding accuracy obtained via multiple- and single-subject feature decoders. The multiple-subject feature decoders were trained on pooled data; the single-subject feature decoders were trained on a single subject’s data. The accuracies were obtained from the source subjects’ test dataset of natural images (error bars, 95% C.I. from 20 individual pairs). (C) Reconstructed natural images for the multiple- and single-subject conditions. (D) Identification accuracies with the natural images. The identification analysis was performed using the pixel values and the extracted DNN feature values of the reconstructions obtained via multiple- and single-subject feature decoders (error bars, 95% C.I. from 20 individual pairs; dotted lines, chance level = 50%).

DNN feature decoding analysis on the natural images showed a small improvement in accuracy across all layers in the multi-subject condition as compared with the single-subject condition. The multiple-subject condition outperformed the single-subject condition both in individual pairs (Fig. S15A) and at the group level (Fig. 7B; ANOVA, effect of decoder type, *F*(1, 361) = 1968, *p* < .001, 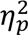 = .85). Similar results were obtained for artificial images, with the multiple-subject condition showing higher accuracies (see Fig. S16A for individual pair results and Fig. S16B for group results; ANOVA, effect of decoder type, *F*(1, 361) = 172, *p* < .001, 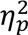 = .32). Reconstructed images obtained using both the single- and multiple-subject decoders showed recognizable visual quality, but the visual qualities were not substantially different (Fig. 7C; see Fig. S16C for artificial images). In the identification analysis of the reconstructed natural images, the multiple-subject condition showed slightly higher accuracies than the single-subject condition (Fig. S15B for individual pairs; ANOVA, effect of decoder type, *p* < .05 in 10/20 individual pairs; Fig. 7D for group results; effect of decoder type, *F*(1, 171) = 75.6, *p* < .001, 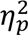 = .30). Similar results were obtained for artificial images, with the multiple-subject condition showing slightly higher identification accuracies than the single-subject condition (Fig. S16D for individual pairs; ANOVA, effect of decoder type, *p* < .05 in 6/20 individual pairs; Fig. S16E for group results; effect of decoder type, *F*(1, 171) = 35.9, *p* < .001, 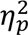 = .17).

We also analyzed the case where there was only limited data for the source subject (300 training samples) to reflect the circumstance where only a small number of data can be collected because of the cost of data collection (Fig. S17). There was also a slight improvement for the identification accuracies of the reconstructed natural images in some of the pairs and at the group level when the multiple-subject feature decoders were used (Fig. S17D for individual pairs; ANOVA, effect of decoder type, *p* < .05 in 5/20 individual pairs; Fig. S17E for group results; effect of decoder type, *F*(1, 171) = 28.3, *p* < .001, 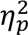 = .14). These results indicate that pooling multiple-subject data is slightly beneficial for improving the accuracy of inter-individual decoding and reconstruction, although the visual quality of the reconstructed images was not largely improved.

We additionally performed different pooling data paradigms by projecting directly other subjects’ data to a subject’s brain space and retraining decoders on the pooled data to examine if this pooling method could achieve any improvement. DNN feature decoding analysis and visual image reconstruction were then performed using the subject’s data. However, the results showed no improvement (Fig. S18, S19).

## 3. Discussion

This study investigated whether and how hierarchical and fine-grained visual information could be converted while preserving perceptual content across individuals using methods of pairwise functional alignment. We first showed that methods of pairwise functional alignment can accurately convert a source subject’s brain activity into a target subject’s brain space by evaluation using the pattern and profile correlations. The ablation analysis on the converters with the exclusion of voxels from various source visual subareas showed that the converters automatically detected the hierarchical correspondences of visual subareas between individuals. We decoded the converted brain activity into DNN features and observed the correspondence between visual subareas and DNN layers. Visual images were reconstructed from the converted brain activity with recognizable shapes and colors of the objects in the presented images. Given sufficient training data, whole VC conversion slightly outperformed the subarea-wise conversion in the inter-individual visual image reconstruction, indicating that the whole VC conversion preserved the hierarchical structure that is explicitly assumed in the subarea-wise conversion. Even with a small number of training data, the converters preserved minimally sufficient information for visual image reconstruction. Pooling data from multiple subjects helped achieve slightly higher accuracy in the visual image reconstruction, even though the visual quality was not greatly improved. Our analyses demonstrate that hierarchically organized fine-grained visual features that enable visual image reconstruction are preserved in the converted brain activity, and functional alignment provide an efficient way to reconstruct visual images without training subject-specific models.

We have shown that the neural code converters automatically detected the hierarchical correspondence of visual subareas between two individuals without explicitly labeling the visual areas (Fig. 2B, C). Previous studies of functional alignment mainly targeted a specific brain area, such as V1 or the inferior temporal cortex (Yamada et al., 2015; Haxby et al., 2011). Other studies functionally aligned a large region of the cortex (Bilenko and Gallant, 2016; Van Uden et al., 2018), but their subsequent analyses targeted different research questions such as the retinotopic organization and the semantic information, leaving the hierarchical correspondences of visual subareas remained undiscussed. Our results explicitly demonstrate that machine learning-based neural code converters can learn the hierarchical correspondence of visual subareas between two individuals. Moreover, some inter-regional predictions existed (*e.g.*, source V1 could predict voxel values in target V2, Fig. 2C), but the prediction accuracy decreases as the cortical distance between two areas increases, presumably indicating that two close areas share some common information.

By decoding the converted fMRI activity patterns into DNN features and reconstructing them as visual images via the decoded DNN features (Fig. 3 and 4), we showed that hierarchically organized fine-grained visual features that enable visual image reconstruction are preserved in the neural code conversion. Previous studies have mainly focused on some specific features, such as object categories, image contrast, retinotopy, and semantics (Haxby et al., 2011; Yamada et al., 2015; Bilenko and Gallant, 2016; Van Uden et al., 2018), but whether a set of hierarchical fine-grained features is preserved after functional alignment remained unknown. The results of DNN feature decoding on multiple levels of DNN layers showed that the converted fMRI activity patterns held multiple levels of fine-grained visual features (Fig. 3). Successful visual image reconstruction further confirmed that the converted fMRI activity patterns preserved sufficient perceptual content to reconstruct visual images (Fig. 4).

The subarea-wise conversion slightly underperformed the whole VC conversion with sufficient training data in the inter-individual visual image reconstruction (Fig. 5 and 6), with the whole VC conversions achieving slightly higher DNN feature decoding accuracy and identification accuracy in the reconstruction. This result shows that the information about explicit labels of visual subareas was not helpful in improving the reconstructed image quality and the whole VC conversion enabled such information to be implicitly learned when sufficient training data are provided.

Training a full visual image reconstruction model requires an fMRI dataset that is costly and takes a long time to collect. In this study, the DNN feature decoders were trained on 6,000 data samples, which took approximately 800 mins of data collection time. In fMRI studies, this long data collection time is impractical for most people. By trading off some visual quality of the reconstructed images, fewer data, for instance, 300 samples, can be collected to train a neural code converter and perform inter-individual visual image reconstruction. In particular, the neural code converter is thought to capture the relationships between individuals’ voxels under a variety of visual scenes, and it could potentially be used together with other decoding models. The inter-individual decoding method with the neural code converter could thus reduce the time and costs of fMRI data collection.

Our results show that pooling data from multiple subjects did not greatly improve the visual image reconstruction (Fig. 7). A possible reason is the variability of data quality, with some subjects’ data leading to relatively poor visual quality in the reconstructed images. Poor quality data limited the capability of the decoders to take advantage of the pooled data, resulting in a limited improvement in visual image reconstruction performance. Furthermore, linear regression models might not be able to fully resolve the feature mismatch of brain activity patterns between individuals, more advanced methods are probably necessitated to solve this problem (Li et al., 2021). Although our analysis showed that pooling data did not lead to great improvements in visual image reconstruction quality, it is still a promising direction for future fMRI studies.

One observation was that the inter-individual image reconstruction did not outperform the within individual image reconstruction (Fig. 4 and 5). As mentioned above, the linear constraint used for conversion might be too restrictive for learning more complex statistical relationships such as nonlinearity between brain activity patterns. Moreover, apart from a consistent stimulus-evoked response across individuals, a brain response to a stimulus also comprises an idiosyncratic stimulus-evoked response and a noise component (Nastase et al., 2019). The brain decoders might leverage the idiosyncratic responses that could not be converted across subjects. Together with the noise components, the inter-individual visual image reconstruction thus underperformed the within-individual visual image reconstruction.

Our results showed that brain activity patterns can be translated across individuals while preserving sufficient information to visualize the perceived stimulus. This provides an efficient way to reconstruct visual images without training subject-specific models, especially when the reconstruction model is complex and data hungry. By reducing the need to collect a large number of data, our approach might prove useful in popularizing the use of brain-machine/computer interfaces that communicate with our internal world.

## 4. Materials and methods

### 4.1 fMRI datasets

#### 4.1.1 Subjects

In this study, Subject 1–3 correspond to the three subjects in Shen et al. (2019a) and the dataset was reused. Subject 4 (male, age 22) and Subject 5 (male, age 27) participated in our additional experiments for the test natural-image and artificial-image sessions. The dataset of the training natural-image session of Subject 4 and 5 was reused from Horikawa and Kamitani (2022). All subjects provided written informed consent for participation in the experiments, in accordance with the Declaration of Helsinki, and the study protocol was approved by the Ethics Committee of Advanced Telecommunications Research Institute International (ATR).

#### 4.1.2 Stimuli

The natural image stimuli in Horikawa and Kamitani (2017) were selected from 200 representative categories in the ImageNet dataset (2011, fall release; Deng et al., 2009). The natural training images were 1,200 images taken from 150 object categories, and the natural test images were 50 images taken from the remaining 50 object categories. The artificial image stimuli used in Shen et al. (2019) consisted of 40 combinations of five shapes (square, small frame, large frame, plus sign, and cross sign) and eight colors (red, green, blue, cyan, magenta, yellow, white, and black).

#### 4.1.3 Experimental design

In both Horikawa and Kamitani (2017), and Shen et. al. (2019), the fMRI signals were measured while subjects viewed sequences of visual images. The visual images had a central fixation spot and were flashed at 2 Hz. Each presentation of an image lasted for 8 s in a stimulus block with four volume scans (Repetition time [TR] = 2 s). The subjects were asked to fixate on the central fixation spot and to click a button when two sequential blocks presented the same image.

The test natural-image session and test artificial-shape session consisted of 24 and 20 runs, respectively. Each run consisted of 55 and 44 stimulus blocks comprising 50 and 40 blocks of different images, and 5 and 4 randomly interspersed repetition blocks, with additional 32-s and 6-s rest periods at the beginning and the end. The 50 natural images and 40 artificial images were presented in random order in each run.

#### 4.1.4 fMRI data preprocessing

The following description is provided by fMRIPrep (https://fmriprep.org/en/1.2.1/citing.html). The results included in this manuscript are based on the data preprocessed using fMRIPrep version 1.2.1 (Esteban et al., 2018) and a Nipype-based tool (Gorgolewski et al., 2011; Gorgolewski et al., 2017). Each T1w (T1-weighted) volume was corrected for INU (intensity non-uniformity) using N4BiasFieldCorrection v2.1.0 (Tustison et al., 2010) and skull-stripped using antsBrainExtraction.sh v2.1.0 (using the OASIS template). Brain surfaces were reconstructed using recon-all from FreeSurfer v5.3.0 (Dale et al., 1999), and the brain mask estimated previously was refined with a custom variation of the method to reconcile ANTs-derived and FreeSurfer-derived segmentations of the cortical gray-matter of Mindboggle (Klein et al. 2017). Spatial normalization to the ICBM 152 Nonlinear Asymmetrical template version 2009c (Fonov et al., 2009) was performed through nonlinear registration with the antsRegistration tool of ANTs v2.1.0 (Avants et al., 2008), using brain-extracted versions of both T1w volume and template. Brain tissue segmentation of cerebrospinal fluid (CSF), white-matter (WM) and gray-matter (GM) was performed on the brain-extracted T1w using fast (Zhang et al., 2001; FSL v5.0.9).

Functional data were slice time corrected using 3dTshift from AFNI v16.2.07 (Cox et al., 1996) and motion corrected using mcflirt (FSL v5.0.9; Jenkinson et al., 2002). This was followed by co-registration to the corresponding T1w using boundary-based registration (Greve et al., 2009) with 9 degrees of freedom, using bbregister (FreeSurfer v6.0.1). Motion correcting transformations, BOLD-to-T1w transformation, and T1w-to-template (MNI) warp were concatenated and applied in a single step using antsApplyTransforms (ANTs v2.1.0) using Lanczos interpolation.

Physiological noise regressors were extracted by applying CompCor (Behzadi et al., 2007). Principal components were estimated for the two CompCor variants: temporal (tCompCor) and anatomical (aCompCor). A mask to exclude signals with cortical origin was obtained by eroding the brain mask, ensuring that it only contained subcortical structures. Six tCompCor components were then calculated including only the top 5% variable voxels within that subcortical mask. For aCompCor, six components were calculated within the intersection of the subcortical mask and the union of CSF and WM masks calculated in T1w space, after their projection to the native space of each functional run. Frame-wise displacement (Power at al., 2013) was calculated for each functional run using the implementation of Nipype.

Many internal operations of fMRIPrep use Nilearn (Abraham et al., 2014), principally within the BOLD-processing workflow. For more details of the pipeline see http://fmriprep.readthedocs.io/en/latest/workflows.html.

The coregistered data to the T1w space were then re-interpolated to 2 × 2 × 2 mm voxels. The data samples were first shifted by 4-s (two volumes) to compensate for the hemodynamic delay, followed by regression to remove nuisance parameters such as a constant baseline, linear trend, and six head motion parameters from each voxel amplitude for each run. The data samples were then despiked to reduce extreme values (beyond ±3 standard deviations for each run) and were averaged within each 8-s trial (four volumes).

### 4.2 Regions of interest (ROIs)

Regions V1, V2, V3, and V4 were delineated using the standard retinotopy experiment (Engel et al., 1994; Sereno et al., 1995) in each subject’s naive brain space. The higher visual cortex (HVC) was defined as a contiguous region covering the LOC, FFA, and PPA, which were identified using conventional functional localizers (Kourtzi and Kanwisher, 2000; Kanwisher et al., 1997; Epstein and Kanwisher, 1998). The whole visual cortex (VC) was defined as the combined regions of V1, V2, V3, V4, and HVC.

### 4.3 Anatomical alignment

For the analyses with anatomical alignment, the subjects’ structural and functional images were nonlinearly normalized to a standard space: the ICBM 152 Nonlinear Asymmetrical template version 2009c (MNI152NLin2009cAsym [MNI space]; see Materials and Methods: “fMRI data preprocessing”). The T1w reference image was spatially normalized to MNI space by the ANTs (Avants et al. 2008) and the functional data were coregistered to this normalized T1w reference image. The coregistered data were then re-interpolated to 2 × 2 × 2 mm voxels. Furthermore, ANTs were used to normalize the ROI masks of V1, V2, V3, V4, and HVC in their native space to the brain in MNI space. In the inter-individual analysis, if a voxel of a source subject and a voxel of a target subject shared the same coordinates, the fMRI activity of the source voxel was considered to be that of the corresponding target voxel. Thus, the voxels of a source subject covered by a ROI mask of a target subject were selected as the input to the model.

### 4.4 Methods of functional alignment

#### 4.4.1 Neural code converter

The neural code converter model for each pair of subjects comprised a set of regularized linear regression models (ridge regression), each trained to predict the activities of an individual voxel of one subject (target) from the brain activity patterns of another subject (source) given the same stimuli. A converter takes a source subject’s brain activity pattern 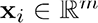 consisting of voxels’ values, and predicts the target brain activity pattern 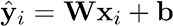, where 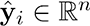 is the converted brain activity pattern consisting of 𝑛 voxels’ values; 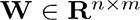 is the conversion matrix and 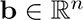 is the bias vector. The converter is trained to minimize the objective function

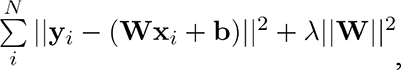

where 𝑦_𝑖_ is the measured target subject’s brain activity pattern for the *i*-th sample, 𝑁 is the number of training samples, λ is the regularization parameter, and 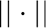 represents the Frobenius norm.

To finetune the regularization parameter λsuch that the performance of visual image reconstruction could be optimized, we performed a 5-fold cross-validation on the training data. At each fold, the brain activity in the validation set was converted to the target’s brain space and then decoded into DNN features. The decoded DNN features were used to calculate an identification accuracy that measured how well a decoded DNN feature pattern can identify the true stimulus between two alternatives (see Materials and Methods: “Identification analysis”). The regularization parameter was optimized in a grid-search manner to maximize the identification accuracy, which is linked to the performance of visual image reconstruction. 500 units instead of all from each layer of the VGG19 model were chosen to save the computational time, and were randomly chosen because there was no a priori knowledge about which DNN units lead to a better visual image reconstruction.

#### 4.4.2 Procrustes transformation

Procrustes transformation is a transformation that includes translation, rotation and uniform scaling, and preserves the shape of a geometric object. It was first introduced by Haxby et al. (2011) in the hyperalignment analysis. Considering the source and target subjects’ brain activity patterns 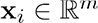 and 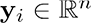, Procrustes transformation estimates an orthogonal transformation matrix 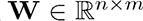 to minimize

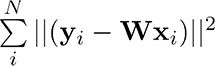

with the constraint 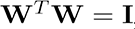 where 𝑁 is the number of training samples. Please refer to Bazeille et al. (2021) for more discussions.

#### 4.4.3 Optimal transport

Optimal transport is related to the question of how one could transform a probability distribution into another probability distribution with the least cost. It was first applied to the functional alignment in Bazeille et al. (2019). Defining 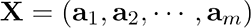 and 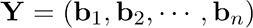 with 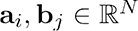 representing a sequence of a voxel response to 𝑁stimuli, optimal transport tries to find a transformation matrix W* such that

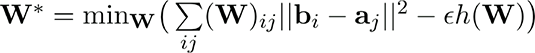

with the constraints

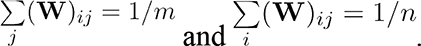

The entropic term

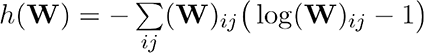

regularizes the optimal transport problem and controls the strength of regularization. The regularization parameter was optimized as in the neural code converter. We used *fmrialign* (https://parietal-inria.github.io/fmralign-docs/index.html) package in the analysis. Please refer to Bazeille et al. (2019) for more mathematical details.

#### 4.4.4 Template-based pairwise alignment via hyperalignment

Template-based pairwise alignment via hyperalignment first estimates a common template among subjects using hyperalignment (Haxby et al., 2011), and then construct a pairwise transformation by first mapping a source subject’s brain activity onto the template, followed by an inverse mapping from the template to a target subject’s brain space. In the first iteration, the hyperalignment algorithm first selects an initial target subject as a template, then aligns the second subject’s fMRI responses to the template using Procrustes transformation.The template is then updated as the mean of the current template and the newly aligned fMRI responses.The same procedure is repeated for additional subjects. In the second iteration, each subject’s original response is aligned to the mean aligned responses of other subjects.The mean aligned response is recalculated and treated as a template. In the last step, each subject’s response is aligned to the template, and an orthogonal transformation matrix is obtained for each subject.

Although the hyperalignment algorithm can estimate the shared space of more than two subjects, we used it only between two subjects, as in the neural code converter analysis.

### 4.5 Noise ceiling estimation

Repeated measures of the brain responses to an identical stimulus are significantly affected by measurement noise in fMRI data, inevitably lowering the prediction accuracy. To account for the noise, we adopted the noise ceiling estimation used by Lescroart and Gallant (2019; see also Hsu et al., 2004). The noise ceiling was obtained by averaging the profile or pattern correlation coefficients between repetitions of the same stimuli within a subject. This noise ceiling estimation is based on the rationale that any model could not predict better than the subject’s own responses. Thus, the noise ceilings reflect the maximum performances of the converter models and were used to normalize the raw prediction accuracies of the converter models (divide raw accuracies by noise ceilings).

Samples/voxels with corresponding noise ceilings below a threshold (99th percentile point in the distribution from random pairs) were excluded from the performance evaluation of the conversion because they could not be reliably measured. But all voxels were included in the downstream DNN feature decoding analysis to prevent information leakage.

### 4.6 DNN model

We used the VGG19 DNN model (Simonyan and Kisserman, 2014) implemented using the Caffe library (Jia et al., 2014). This model is pre-trained for the 1,000-class object recognition task using the images from ImageNet (Deng et al., 2009; the pre-trained model is available from https://github.com/BVLC/caffe/wiki/Model-Zoo). The model consists of 16 convolutional layers and three fully connected layers. All the input images to the model were rescaled to 224 ྾ 224 pixels. Following Shen et al. (2019), outputs from individual units before rectification were used as target variables in the DNN feature decoding analysis. The number of units in each layer is as follows: conv1_1 and conv1_2, 3,211,264; conv2_1 and conv2_2, 1,605,632; conv3_1, conv3_2, conv3_3, and conv3_4, 802,816; conv4_1, conv4_2, conv4_3, and conv4_4, 401,408; conv5_1, conv5_2, conv5_3, and conv5_4, 100352; fc6 and fc7, 4,096; and fc8, 1,000.

We used the AlexNet DNN model (Krizhevsky et al., 2012) implemented using the Caffe library to extract DNN features from the reconstructed images and the presented image. This model is also pre-trained similarly (available from https://github.com/BVLC/caffe/tree/master/models/bvlc_alexnet). The model consists of five convolutional layers and three fully connected layers. The number of units in each layer is as follows: conv1, 290,400; conv2, 186,624; conv3 and conv4, 64,896; conv5, 43,264; fc6 and fc7, 4,096; and fc8, 1,000.

### 4.7 DNN feature decoding analysis

For each DNN unit, we trained a ridge linear regression model (DNN feature decoder) that takes an fMRI activity pattern induced by a stimulus as input and predicts a feature value of the stimulus. The ridge regularization parameter was set to 100. Before model training, the feature values and the voxel values were normalized, and then a voxel selection procedure was performed: the Pearson correlation coefficients between the sequences of feature values and voxel responses of all voxels were computed and the top 500 voxels having the highest correlations were selected for training. The trained decoders were tested on the average fMRI pattern over repetitions to increase the signal-to-noise ratio of the fMRI signal. For details of the feature decoding, see Horikawa and Kamitani (2017, 2022) and Shen et al. (2019a).

### 4.8 Brain hierarchy (BH) score

The BH score was originally designed to measure the degree to which an artificial neural network is hierarchically similar to the human brain (Nonaka et al., 2021). In this study, we adopted the decoding-based BH score to see whether the hierarchical similarity is preserved after inter-individual conversion. The DNN features of randomly selected 1,000 units of each layer are decoded from the fMRI pattern of one of the five visual areas: V1–V4 and the HVC. For each unit, the visual area showing the best decoding accuracy was identified and was called the “top visual area.” The first layer, the last layer, and three randomly sampled intermediate layers were used to calculate a Spearman rank correlation coefficient between the hierarchical levels of the five DNN layers (coded as 0 through 4) and the top visual area (coded as V1: 0, V2: 1, V3: 2, V4: 3, and HVC: 4) across DNN units. This sampling procedure was repeated 10,000 times, and the mean Spearman rank correlation coefficient was taken as the BH score. See Nonaka et al. (2021) for more details.

### 4.9 Visual image reconstruction

We used an image reconstruction method (deep image reconstruction) proposed by Shen et al. (2019). The method optimizes pixel values of an input image based on a set of DNN features given as a target. Given the decoded DNN features from multiple layers, an image was reconstructed by solving the following optimization problem (Mahendran et al., 2015):

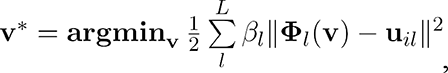

Where 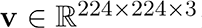 is a vector whose elements are the pixel values of an image (width ྾ height ྾ RGB channels); 𝐿 is the total number of layers;Φ_𝑙_ is the function that maps the image to the DNN feature vector of the 𝑙-th layer; u_𝑙𝑖_ is the decoded DNN feature vector of the 𝑙-th layer for the i-th sample; and 𝛽_𝑙_ is the parameter that weights the contribution of the 𝑙-th layer, which was set to be 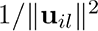.

A natural image prior is applied by introducing a generative adversarial network called the deep generator network (DGN) to enhance the naturalness of the image (Nguyen et al., 2016). The optimization problem becomes

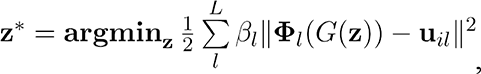

where 𝐺 is the DGN and z is a latent vector. The reconstructed image is obtained by 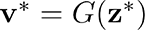. The DGN is a pre-trained generator provided by Dosovitskiy and Brox (2016; available from https://github.com/dosovits/caffe-fr-chairs).

The solution to the above optimization problem is considered to be the reconstructed image from the brain activity pattern. Following Shen et al. (2019a), natural images were reconstructed using the DGN, and the objective function was optimized by stochastic gradient descent with momentum with 200 iterations, whereas the artificial images were reconstructed without the DGN and the objective function was optimized by a limited-memory BFGS algorithm with 200 iterations (Le et al., 2011; Liu and Nocedal, 1989; Gatys et al., 2016).

### 4.10 Identification analysis

Identification analysis was used to evaluate image reconstruction quality. Presented images were identified using the similarity in either image pixels or DNN features, which were reshaped into a one-dimensional feature vector. The feature vector of a reconstructed image was used to compare the true feature vector of the presented image with the false alternative of another image. The comparison was counted as correctly identified if the feature vector of the reconstruction has a higher Pearson correlation coefficient with the true feature vector than with the false alternative. The identification was repeated for multiple false alternatives for each reconstruction. For natural images, the identification was repeated with 49 alternatives for each reconstruction, resulting in 50 images × 49 comparisons = 2,450 comparisons in total. The identification accuracy for a reconstructed image was defined as the proportion of correct identification.

When optimizing the regularization parameters for the neural code converters by cross validation, we used a set of decoded DNN features concatenated from multiple layers to calculate the identification accuracies to evaluate the performance (see Materials and Methods: “Methods of functional alignment”). The candidate images for comparisons were a subset of the 1,200 images presented in the training image session.

### 4.11 Statistics

We performed statistical analyses primarily on the data samples in each pair of subjects to examine the effect of each conversion and the prevalence across pairs (Ince et al., 2022). We additionally performed group level analyses using the mean values from 20 individual pairs for summary purposes or for cases where within-pair analysis was not applicable. In some analyses, the results with converted brain activity were compared with those without conversion (within-individual), in which the data from five subjects were similarly processed.

In the evaluation of conversion for each pair of subjects, the pattern correlation coefficients for 50 visual stimuli were used to calculate the mean conversion accuracy (pattern) and its 95% confidence interval, while the profile correlation coefficients for all voxels were used to calculate the mean conversion accuracy (profile) and its 95% confidence interval. At the group level, the mean conversion accuracies (pattern/profile) from 20 individual pairs were used to calculate the group mean and its 95% confidence interval.

In the DNN feature decoding analysis for each conversion or each subject (within), the decoding accuracies (profile correlations in individual units) for all DNN units were used to calculate the mean decoding accuracy and its 95% confidence interval. At the group level, the mean decoding accuracies from 20 individual pairs or five subjects (within) were used to calculate the group mean and its 95% confidence interval.

In the evaluation of the brain hierarchy, a BH score was computed for each conversion or each subject (within). At the group level, the BH scores of 20 individual pairs or five subjects (within) were averaged to obtain the mean BH score.

In the identification analysis of reconstructed images for each conversion or each subject (within), the identification accuracies for individual reconstructed images were used to calculate the mean identification accuracy and its 95% confidence interval. At the group level, the mean identification accuracies from 20 individual pairs or five subjects (within) were used to calculate the group mean and its 95% confidence interval.

In the comparison of the subarea-wise conversion with the whole VC conversion, we performed ANOVA on DNN feature decoding accuracies and identification accuracies with the conversion type as a repeated measure factor and the DNN layer as a between-subject factor. Since millions of DNN units and their decoding accuracies (profile correlations) always lead to statistical significance, we only performed group level ANOVA on DNN feature decoding accuracies to compute the *F* scores, *p* values, and the effect size, in which the data points were the mean DNN feature decoding accuracies of 20 individual pairs. For identification, the accuracies with individual reconstructed images were used as the data points in the ANOVA analysis to compute the *F* scores, *p* values, and the effect size in each individual pair. At the group level, the data points were the mean identification accuracies from 20 individual pairs, which were used to compute the *F* scores, *p* values, and the effect size.

Similar ANOVA tests were applied to the pooling data analysis to examine the effect of decoder type (multiple- and single-subject).

## Data and code availability

The experimental code and data that support the findings of this study are respectively available from our repository (code for inter-individual deep image reconstruction including neural code converter, Procrustes transformation, optimal transport and hyperalignment: https://github.com/KamitaniLab/InterIndividualDeepImageReconstruction, code for feature decoding: https://github.com/KamitaniLab/dnn-feature-decoding, code for image reconstruction: https://github.com/KamitaniLab/DeepImageReconstruction, code for BH score calculation: https://github.com/KamitaniLab/BHscore) and open data repository (OpenNeuro: https://openneuro.org/datasets/ds003993/versions/1.0.0 for Subject 1–3, https://openneuro.org/datasets/ds003430/versions/1.2.0 for the dataset of training natural-image session for Subject 4 and 5, and https://openneuro.org/datasets/ds001506/versions/1.3.1 for the dataset of test natural-image and artificial-image sessions for Subject 4 and 5).

## Author contributions

**Jun Kai Ho**: Methodology, Software, Formal analysis, Data curation, Writing - original draft, Writing - review & editing, Visualization; **Tomoyasu Horikawa**: Methodology, Validation, Investigation, Writing - original draft, Writing - review & editing; **Kei Majima**: Methodology, Writing - review & editing; **Fan Cheng**: Methodology; **Yukiyasu Kamitani**: Conceptualization, Validation, Investigation, Resources, Writing - original draft, Writing - review & editing, Supervision, Project administration, Funding acquisition.

## Ethics statement

All subjects provided written informed consent for participation in the experiments, in accordance with the Declaration of Helsinki, and the study protocol was approved by the Ethics Committee of Advanced Telecommunications Research Institute International (ATR).

## Acknowledgments

The authors would like to thank Jong-Yun Park, Shangfeng Jin, Shuntaro C. Aoki, Mitsuaki Tsukamoto, and Misato Tanaka for helpful comments on the manuscript. The data used in the study were collected using the fMRI scanner and related facilities of Kokoro Research Center, Kyoto University. We thank Kimberly Moravec, PhD, from Edanz (https://jp.edanz.com/ac) for editing a draft of this manuscript.

## References

1. Abraham A, Pedregosa F, Eickenberg M, Gervais P, Mueller A, Kossaifi J, et al. Machine learning for neuroimaging with scikit-learn. Front in Neuroinf. 2014;8:14. doi: 10.3389/fninf.2014.00014.

2. Avants BB, Epstein CL, Grossman M, Gee JC. Symmetric diffeomorphic image registration with cross-correlation: Evaluating automated labeling of elderly and neurodegenerative brain. Medical Image Analysis. 2008;12: 26–41. doi: 10.1016/j.media.2007.06.004.

3. Bazeille T, DuPre E, Richard H, Poline JB, Thirion B. An empirical evaluation of functional alignment using inter-subject decoding. NeuroImage. 2021;245: 118683. doi: 10.1016/j.neuroimage.2021.118683

4. Bazeille T, Richard H, Janati H, Thirion B. Local Optimal Transport for Functional Brain Template Estimation. Information Processing in Medical Imaging; June 2019; Hong Kong. Springer, Cham; vol 11492. doi: 10.1007/978-3-030-20351-1_18

5. Behzadi Y, Restom K, Liau J, Liu TT. A component based noise correction method (CompCor) for BOLD and perfusion based fMRI. Neuroimage. 2007;37:90–101. doi: 10.1016/j.neuroimage.2007.04.042.

6. Bilenko NY, Gallant JL. Pyrcca: Regularized kernel canonical correlation analysis in Python and its applications to neuroimaging. Front Neuroinform. 2016;10: 49. doi: 10.3389/fninf.2016.00049.

7. Blumensath T, Jbabdi S, Glasser MF, Van Essen DC, Ugurbil K, Behrens TEJ, et al. Spatially constrained hierarchical parcellation of the brain with resting-state fMRI. NeuroImage. 2013;76: 313–324. doi: 10.1016/j.neuroimage.2013.03.024.

8. Chen P-H, Chen J, Yeshurun Y, Hasson U, Haxby J, Ramadge PJ. A reduced-dimension fMRI shared response model. Adv Neural Inf Process Syst. 2015;28: 460–468. Available from: http://papers.nips.cc/paper/5855-a-reduced-dimension-fmri-shared-response-model.pdf

9. Cox RW. AFNI: software for analysis and visualization of functional magnetic resonance neuroimages. Comput Biomed Res. 1996;29: 162–173. doi: 10.1006/cbmr.1996.0014.

10. Dale A, Fischl B, Sereno MI. Cortical Surface-Based Analysis: I. Segmentation and Surface Reconstruction. Neuroimage. 1999;9: 179–194. doi: 10.1006/nimg.1998.0395.

11. Deng J, Dong W, Socher R, Li L, Li K, Fei-Fei L. ImageNet: A large-scale hierarchical image database. Proceedings of the IEEE Computer Society Conference on Computer Vision and Pattern Recognition; June 2009; Miami, FL, USA. IEEE; 2009. p. 248–255. doi: 10.1109/CVPR.2009.5206848.

12. Dosovitskiy A, Brox T. Generating images with perceptual similarity metrics based on deep networks. Adv Neural Inf Process Syst. 2016;29: 658–666. Available from: https://arxiv.org/abs/1602.02644

13. Engel SA, Rumelhart DE, Wandell BA, Lee AT, Glover GH, Chichilnisky E-J, et al. fMRI of human visual cortex. Nature. 1994;369: 525. doi: 10.1038/369525a0.

14. Epstein R, Kanwisher N. A cortical representation of the local visual environment. Nature. 1998;392: 598–601. doi: 10.1038/33402.

15. Esteban O, Markiewicz CJ, Blair RW, Moodie CA, Isik AI, Erramuzpe A, et al. fMRIPrep: A robust preprocessing pipeline for functional MRI. Nature Methods. 2019;16: 111–116. doi: 10.1038/s41592-018-0235-4.

16. Fischl B, Rajendran N, Busa E, Augustinack J, Hinds O, Yeo BTT, et al. Cortical folding patterns and predicting cytoarchitecture. Cerebral Cortex. 2008;18: 1973–1980. doi: 10.1093/cercor/bhm225.

17. Fonov VS, Evans AC, McKinstry RC, Almli CR, Collins DL. Unbiased nonlinear average age-appropriate brain templates from birth to adulthood. NeuroImage; Amsterdam. 2009;47: S102. doi: 10.1016/S1053-8119(09)70884-5.

18. Gatys LA, Ecker AS, Bethge M. Image style transfer using convolutional neural networks. Proceedings of the IEEE Computer Society Conference on Computer Vision and Pattern Recognition. December 2016; Las Vegas, NV, USA. IEEE; 2016. p. 2414–2423. doi: 10.1109/CVPR.2016.265.

19. Gorgolewski K, Burns CD, Madison C, Clark D, Halchenko YO, Waskom ML, et al. Nipype: a flexible, lightweight and extensible neuroimaging data processing framework in python. Front Neuroinform. 2011;5: 13. doi: 10.3389/fninf.2011.00013.

20. Gorgolewski KJ, Esteban O, Ellis DG, Notter MP, Ziegler E, Johnson H, et al. Nipype: a flexible, lightweight and extensible neuroimaging data processing framework in Python. 2017. doi: 10.5281/zenodo.581704.

21. Greve DN, Fischl B. Accurate and robust brain image alignment using boundary-based registration. Neuroimage. 2009;48: 63–72. doi: 10.1016/j.neuroimage.2009.06.060.

22. Güçlü U, van Gerven MAJ. Deep neural networks reveal a gradient in the complexity of neural representations across the ventral stream. J Neurosci. 2015;35: 10005–10014. doi: 10.1523/JNEUROSCI.5023-14.2015.

23. Güçlü U, van Gerven MAJ. Increasingly complex representations of natural movies across the dorsal stream are shared between subjects. NeuroImage. 2017;145: 329–336. doi: 10.1016/j.neuroimage.2015.12.036.

24. Guntupalli JS, Hanke M, Halchenko YO, Connolly AC, Ramadge PJ, Haxby JV. A model of representational spaces in human cortex. Cerebral Cortex. 2016;26: 2919–2934. doi: 10.1093/cercor/bhw068.

25. Haxby JV, Guntupalli JS, Connolly AC, Halchenko YO, Conroy BR, Gobbini MI, et al. A common, high-dimensional model of the representational space in human ventral temporal cortex. Neuron. 2011;72: 404–416. doi: 10.1016/j.neuron.2011.08.026.

26. Horikawa T, Kamitani Y. Attention modulates neural representation to render reconstructions according to subjective appearance. Commun Biol. 2022;5: 1–12. doi: 10.1038/s42003-021-02975-5.

27. Horikawa T, Kamitani Y. Generic decoding of seen and imagined objects using hierarchical visual features. Nature Communications. 2017;8: 15037. doi: 10.1038/ncomms15037.

28. Hsu A, Borst A, Theunissen FE. Quantifying variability in neural responses and its application for the validation of model predictions. Network: Computation in Neural Systems. 2004;15: 91–109. doi: 10.1088/0954-898X_15_2_002.

29. Hubel DH, Wiesel TN. Receptive fields, binocular interaction and functional architecture in the cat’s visual cortex. The Journal of Physiology. 1962;160: 106–154. doi: 10.1113/jphysiol.1962.sp006837.

30. Ince RAA, Kay JW, Schyns PG. Within-participant statistics for cognitive science. Trends in Cognitive Sciences. 2022;26: 626–630. doi: 10.1016/j.tics.2022.05.008.

31. Jenkinson M, Bannister P, Brady M, Smith S. Improved optimization for the robust and accurate linear registration and motion correction of brain images. Neuroimage. 2002;17: 825–841. doi: 10.1006/nimg.2002.1132.

32. Jia Y, Shelhamer E, Donahue J, Karayev S, Long J, Girshick R, et al. Caffe: Convolutional architecture for fast feature embedding. arXiv:1408.5093 [Preprint]. 2014 [cited 2021 Nov 8]. Available from: https://arxiv.org/abs/1408.5093.

33. Kanwisher N, McDermott J, Chun MM. The fusiform face area: A module in human extrastriate cortex specialized for face perception. J Neurosci. 1997;17: 4302–4311. doi: 10.1523/JNEUROSCI.17-11-04302.1997.

34. Klein A, Ghosh SS, Bao FS, Giard J, Häme Y, Stavsky E, et al. Mindboggling morphometry of human brains. PLoS Comput Biol. 2017;13: e1005350. doi: 10.1371/journal.pcbi.1005350.

35. Kourtzi Z, Kanwisher N. Cortical regions involved in perceiving object shape. J Neurosci. 2000;20: 3310–8. doi: 10.1523/JNEUROSCI.20-09-03310.2000.

36. Krizhevsky A, Sutskever I, Hinton GE. ImageNet classification with deep convolutional neural networks. Adv Neural Inf Process Syst. 2012;25: 1106–1114. Available from: https://papers.nips.cc/paper/2012/hash/c399862d3b9d6b76c8436e924a68c45b-Abstract.html

37. Laumann TO, Gordon EM, Adeyemo B, Snyder AZ, Joo SJ, Chen M-Y, et al. Functional system and areal organization of a highly sampled individual human brain. Neuron. 2015;87: 657–670. doi: 10.1016/j.neuron.2015.06.037.

38. Le QV, Ngiam J, Coates A, Lahiri A, Prochnow B, Ng AY. On optimization methods for deep learning. Proceedings of the 28th International Conference on International Conference on Machine Learning; June 2011; Bellevue, Washington, USA. Omnipress. 2011. p. 265–272.

39. Lescroart MD, Gallant JL. Human scene-selective areas represent 3D configurations of surfaces. Neuron. 2019;101: 178–192. doi: 10.1016/j.neuron.2018.11.004.

40. Li D, Du C, Wang S, Wang H, He H. Multi-subject data augmentation for target subject semantic decoding with deep multi-view adversarial learning. Information Sciences. 2021;547: 1025–44. doi: 10.1016/j.ins.2020.09.012

41. Liu DC, Nocedal J. On the limited memory BFGS method for large scale optimization. Math Program. 1989;45: 503–528. doi: 10.1007/BF01589116.

42. Mahendran A, Vedaldi A. Understanding deep image representations by inverting them. Proceedings of the IEEE Computer Society Conference on Computer Vision and Pattern Recognition; June 2015; Boston, MA, USA. IEEE; 2015 p. 5188–5196. doi: 10.1109/CVPR.2015.7299155.

43. Mishkin M, Ungerleider LG. Contribution of striate inputs to the visuospatial functions of parieto-preoccipital cortex in monkeys. Behavioural Brain Research. 1982;6: 57–77. doi: 10.1016/0166-4328(82)90081-X.

44. Nastase SA, Gazzola V, Hasson U, Keysers C. Measuring shared responses across subjects using intersubject correlation. Soc Cogn Affect Neurosci. 2019;14: 667–685. doi: 10.1093/scan/nsz037.

45. Nguyen A, Dosovitskiy A, Yosinski J, Brox T, Clune J. Synthesizing the preferred inputs for neurons in neural networks via deep generator networks. Adv Neural Inf Process Syst. 2016;29: 3387–3395. Available from: https://arxiv.org/abs/1605.09304

46. Nonaka S, Majima K, Aoki SC, Kamitani Y. Brain hierarchy score: Which deep neural networks are hierarchically brain-like? iScience. 2021;24. doi: 10.1016/j.isci.2021.103013.

47. Power JD, Mitra A, Laumann TO, Snyder AZ, Schlaggar BL, Petersen SE. Methods to detect, characterize, and remove motion artifact in resting state fMRI. Neuroimage. 2013;84: 320–341. doi: 10.1016/j.neuroimage.2013.08.048.

48. Schönemann PH. A generalized solution of the orthogonal procrustes problem. Psychometrika. 1966;31: 1–10. doi: 10.1007/BF02289451.

49. Sereno MI, Dale AM, Reppas JB, Kwong KK, Belliveau JW, Brady TJ, et al. Borders of multiple visual areas in humans revealed by functional magnetic resonance imaging. Science. 1995;268: 889–893. doi: 10.1126/science.7754376.

50. Shen G, Dwivedi K, Majima K, Horikawa T, Kamitani Y. End-to-end deep image reconstruction from human brain activity. Front Comput Neurosci. 2019;13: 21. doi: 10.3389/fncom.2019.00021.

51. Shen G, Horikawa T, Majima K, Kamitani Y. Deep image reconstruction from human brain activity. PLoS Computational Biology. 2019;15: 1006633. doi: 10.1371/journal.pcbi.1006633.

52. Simonyan K, Zisserman A. Very deep convolutional networks for large-scale image recognition. arXiv:1409.1556v1 [Preprint]. 2014 [cited 2021 Nov 8] Available from: https://arxiv.org/abs/1409.1556.

53. Smith PL, Little DR. Small is beautiful: In defense of the small-N design. Psychonomic Bulletin & Review. 2018;25: 2083–101. doi: 10.3758/s13423-018-1451-8

54. Tustison NJ, Avants BB, Cook PA, Zheng Y, Egan A, Yushkevich PA, et al. N4ITK: improved N3 bias correction. IEEE Trans Med Imaging. 2010;29: 1310–1320. doi: 10.1109/TMI.2010.2046908.

55. Van Essen DC. A population-average, landmark- and surface-based (PALS) atlas of human cerebral cortex. NeuroImage. 2005;28: 635–662. doi: 10.1016/j.neuroimage.2005.06.058.

56. Van Essen DC. Surface-based approaches to spatial localization and registration in primate cerebral cortex. NeuroImage. 2004;23: S97–107. doi: 10.1016/j.neuroimage.2004.07.024.

57. Van Uden CE, Nastase SA, Connolly AC, Ma FL., Hansen I, Gobbini MI et al. Modeling semantic encoding in a common neural representational space. Front Neurosci. 2018;12: 437. doi: 10.3389/fnins.2018.00437.

58. Watson JDG, Myers R, Frackowiak RSJ, Hajnal JV, Woods RP, Mazziotta JC, et al. Area V5 of the human brain: Evidence from a combined study using positron emission tomography and magnetic resonance imaging. Cerebral Cortex. 1993;3: 79–94. doi: 10.1093/cercor/3.2.79.

59. Yamada K, Miyawaki Y, Kamitani Y. Inter-subject neural code converter for visual image representation. NeuroImage. 2015;113: 289–297. doi: 10.1016/j.neuroimage.2015.03.059.

60. Yamada K, Miyawaki Y, Kamitani Y. Neural Code Converter for Visual Image Representation. International Workshop on Pattern Recognition in NeuroImaging. May 2011. Seoul, South Korea. IEEE; p. 37–40. doi: 10.1109/PRNI.2011.13.

61. Yamins DLK, Hong H, Cadieu CF, Solomon EA, Seibert D, DiCarlo JJ. Performance-optimized hierarchical models predict neural responses in higher visual cortex. Proc Natl Acad Sci USA. 2014;111: 8619–8624. doi: 10.1073/pnas.1403112111.

62. Zhang Y, Brady M, Smith S. Segmentation of brain MR images through a hidden Markov random field model and the expectation-maximization algorithm. IEEE Trans Med Imaging. 2001;20: 45–57. doi: 10.1109/42.906424.

