## Supplementary materials for "Inter-individual deep image reconstruction"

### Supplementary figures

Fig. S1. Illustration of pairwise alignment and template-based pairwise alignment.

Fig. S2. Conversion accuracy and visualization of brain activity patterns.

Fig. S3. Evaluation of different methods of functional alignment on visual areas.

Fig. S4. The effects of source area exclusion on neural code conversions.

Fig. S5. DNN feature decoding and hierarchical representation of representative individual pairs.

Fig. S6. Inter-individual DNN feature decoding accuracies of different methods of functional alignment.

Fig. S7. Reconstructed natural images across individual pairs.

Fig. S8. Reconstructed artificial images across individual pairs.

Fig. S9. Reconstructed images obtained via different methods of functional alignment.

Fig. S10. Reconstructed images using different repetitions of samples.

Fig. S11. Identification accuracies of representative individual pairs.

Fig. S12. Evaluations of subarea-wise neural code converters.

Fig. S13. Effect of the number of training data for the converter on the reconstruction of artificial images.

Fig. S14. Identification accuracies of representative individual pairs with different numbers of training samples for converters.

Fig. S15. Feature decoding and identification analyses of representative individual pairs with multiple- and single-subject feature decoders.

Fig. S16. Evaluation of multiple subject feature decoders using the artificial images dataset.

Fig. S17. Pooling data analysis with limited data for a novel subject.

Fig. S18. Augmenting data of a subject.

Fig. S19. Augmenting data of a subject with stimulus variation.

Video S1. Deep image reconstruction in within-individual and across-individual conditions.

Video S2. Deep image reconstruction via converters trained with varying numbers of data.

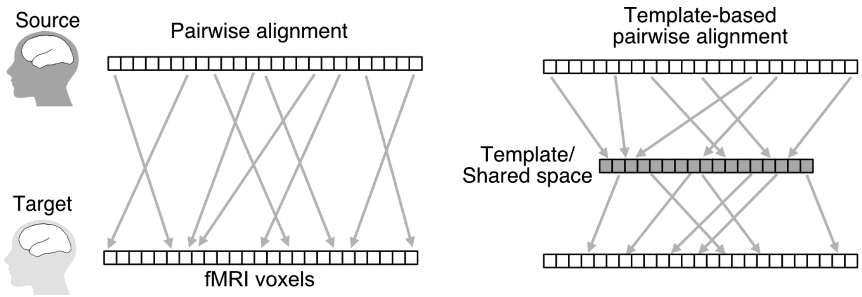


**Fig. S1. Illustration of pairwise alignment and template-based pairwise alignment.** Ridge-based neural code converter, Procrustes transformation and optimal transport are pairwise alignment, and hyperalignment was used to construct a template to conduct template-based pairwise alignment. Analyses for all methods were performed with 2,400 training samples from a pair of source and target subjects. Pairwise alignment directly aligned the source subject's responses to the target subject brain space (left). Template-based pairwise alignment first mapped a source subject's responses into a template, followed by an inverse mapping into the target subject's brain space (right).

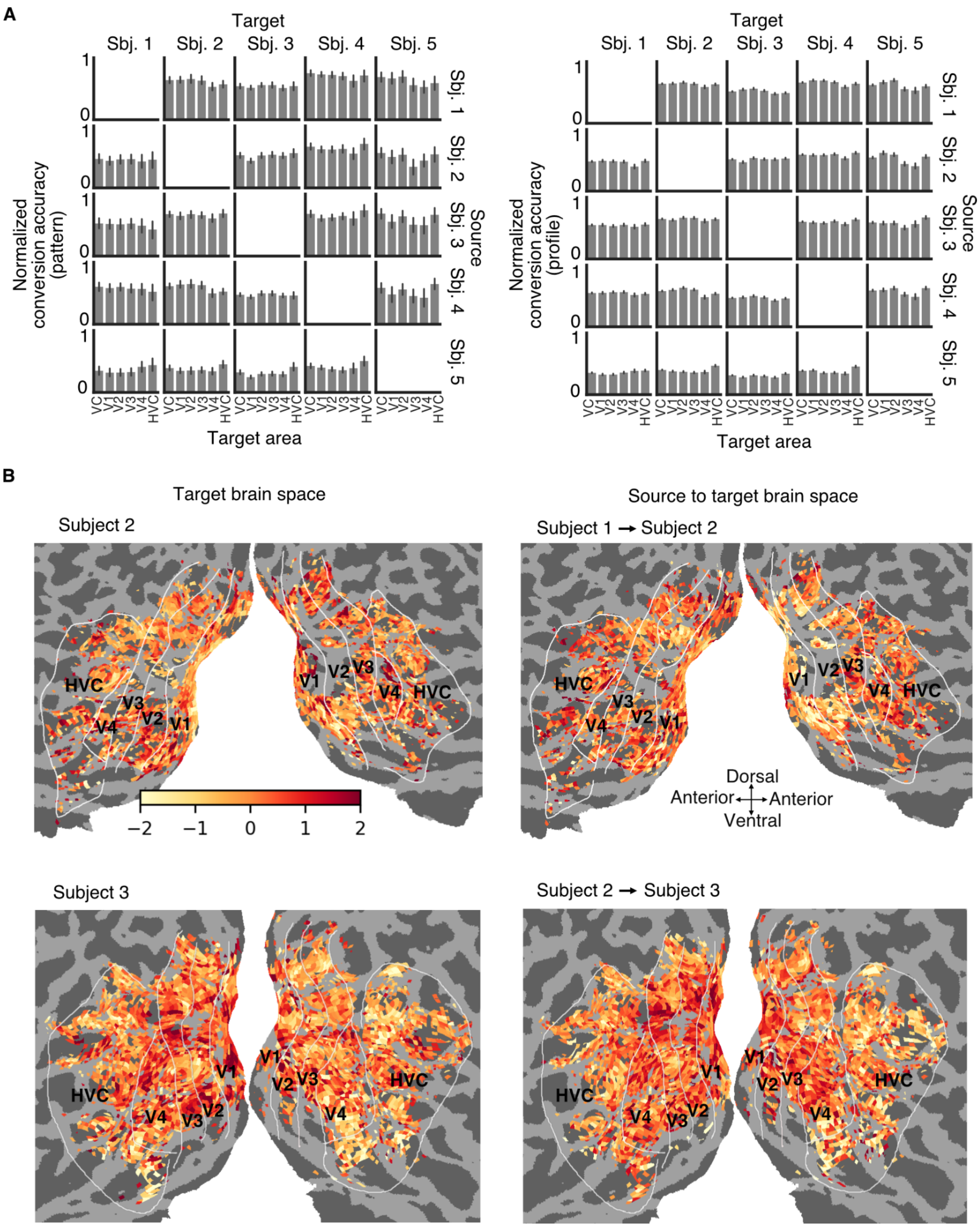


**Fig. S2. Conversion accuracy and visualization of brain activity patterns.**
(A) Conversion accuracies of individual pairs. The pattern correlation coefficients for 50 visual stimuli were used to calculate a mean conversion accuracy (pattern) and its 95% confidence interval. The profile correlation coefficients for voxels were used to calculate a mean conversion

accuracy (profile) and its 95% confidence interval (right; error bars on left panel, 95% C.I. across visual images; error bars on the right panel, 95% C.I. across voxels).

(B) The single trial brain activity patterns responding to two test natural images are plotted. The left panels show the brain activity pattern for the golden fish in Subject 2 and the brain activity pattern for the butterfly in Subject 3 in their native brain space. The right panels show the brain activity patterns converted from the source subjects (Subject 1 and 2) to the target subjects. The activation values were normalized for ease of visualization.

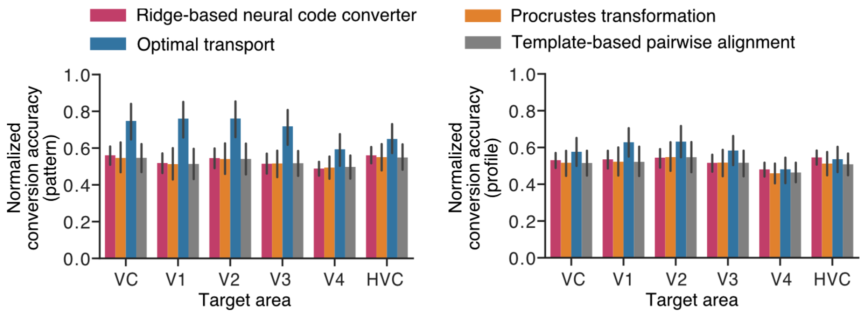


**Fig. S3. Evaluation of different methods of functional alignment on visual areas.**

The conversion accuracy was averaged across 20 individual pairs for Ridge-based neural code converter, Procrustes transformation, optimal transport, and template-based pairwise alignment via hyperalignment (*c.f.* Fig 2B; error bars, 95% confidence interval [C.I.] from 20 individual pairs).

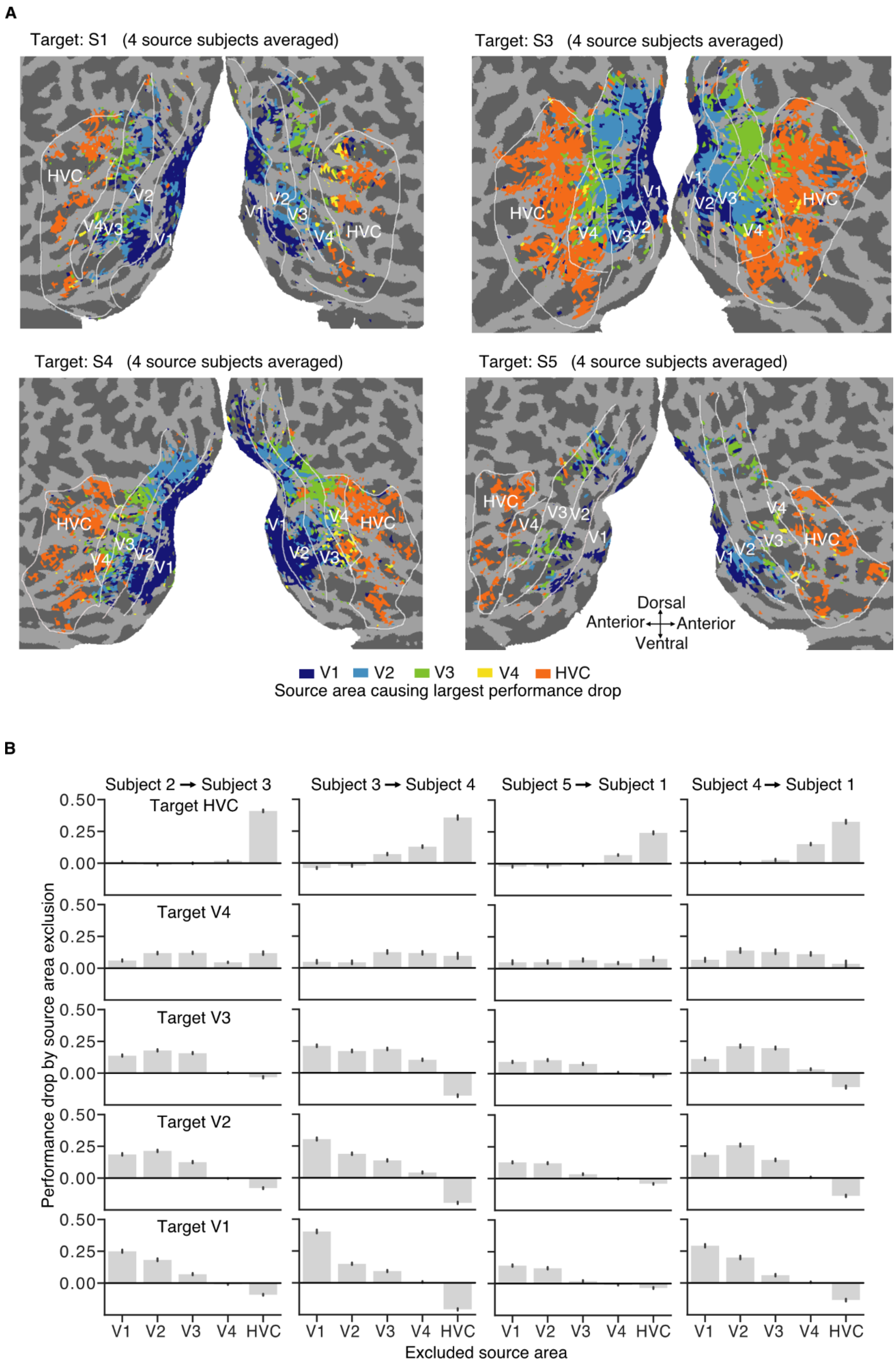
A
Target: S1 (4 source subjects averaged)
Target: S3 (4 source subjects averaged)
Target: S4 (4 source subjects averaged)
Target: S5 (4 source subjects averaged)
HVC
V1
V2
V3
V4
Dorsal
Anterior
Anterior
Ventral
V1 V2 V3 V4 HVC
Source area causing largest performance drop
B
Subject 2 → Subject 3
Subject 3 → Subject 4
Subject 5 → Subject 1
Subject 4 → Subject 1
Target HVC
Target V4
Target V3
Target V2
Target V1
0.50
0.25
0.00
Performance drop by source area exclusion
V1 V2 V3 V4 HVC
Excluded source area

**Fig. S4. The effects of source area exclusion on neural code conversions.**

(A) The cortical map of the effects of source area exclusion. Each voxel on the target brain is colored by the index of the excluded visual area that caused the largest performance drop when the converter models were tested with the test natural image dataset (performance drops were averaged across four source subjects for a single target subject). Only voxels that generate reliable responses with noise ceilings above a threshold are shown (see Materials and Methods: "Noise ceiling estimation"). The target Subject 1, 3, 4, and 5 are shown.

(B) Performance drop caused by source area exclusion for four representative individual pairs. The performance drops for voxels were used to obtain a mean performance drop and its 95% confidence interval (error bar, 95% C.I. of performance drops across voxels).

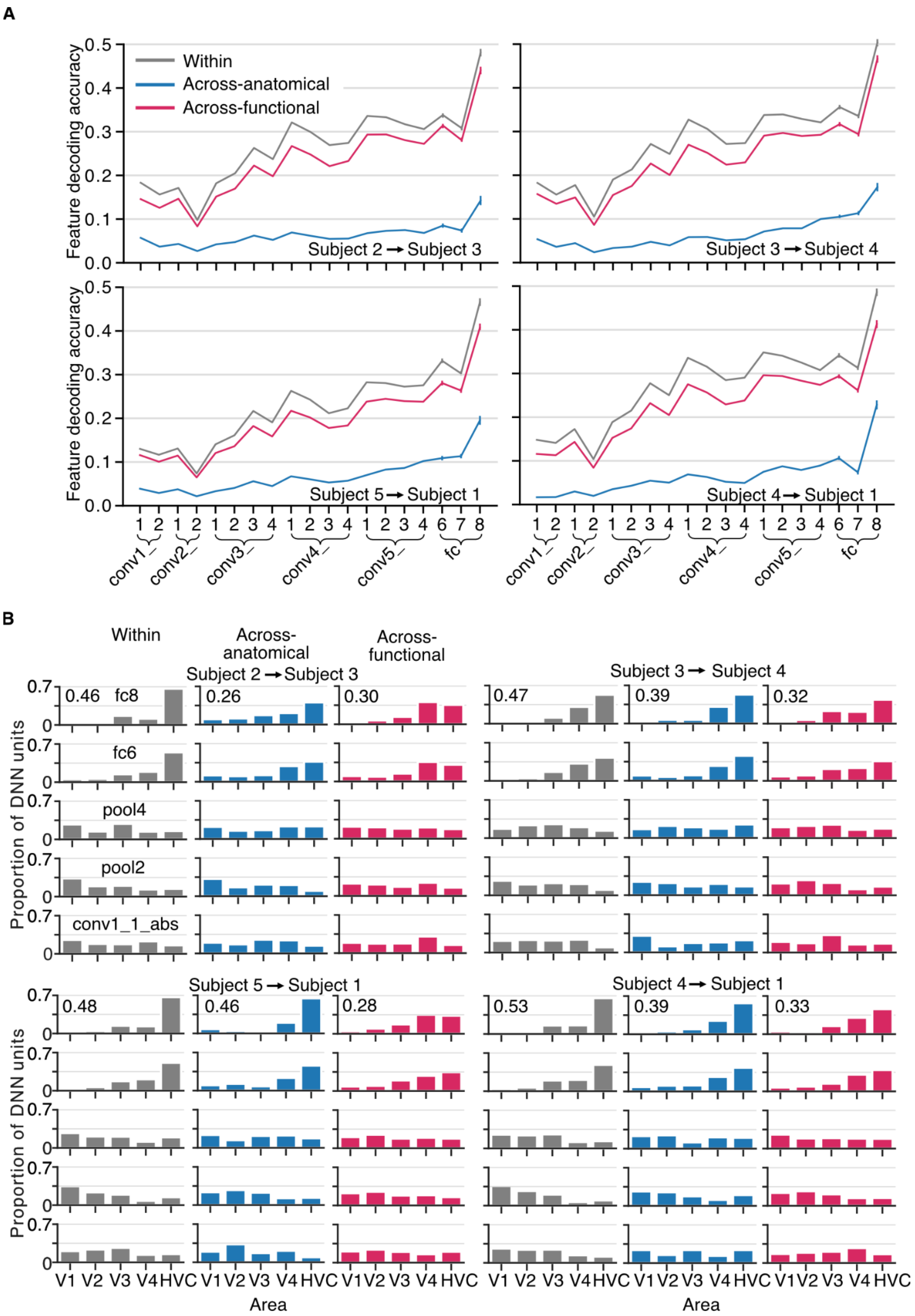


**Fig. S5. DNN feature decoding and hierarchical representation of representative individual pairs.**

(A) DNN feature decoding accuracy from the whole visual cortex (VC). Decoding accuracies for each layer of the VGG19 model are shown for the Within, Across-anatomical, and Across-functional conditions. The decoding accuracies for all DNN units in each layer were used to calculate a mean decoding accuracy and its 95% confidence interval. Results of four representative individual pairs are shown (error bar, 95% C.I. across voxels).
(B) Proportion the "top visual area" (best decodable area for each DNN unit) across DNN units in each layer. Only five representative layers are shown. Each bar indicates the proportion of DNN units. The numbers on the top left indicate the BH scores. Results of four representative individual pairs are shown.

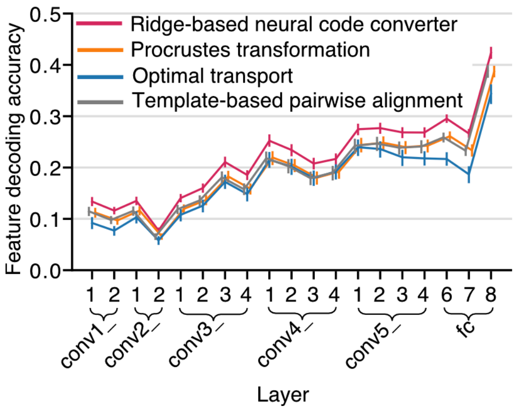


**Fig. S6. Inter-individual DNN feature decoding accuracies of different methods of functional alignment.**

The decoding accuracies were averaged across 20 individual pairs for Ridge-based neural code converter, Procrustes transformation, optimal transport, and template-based pairwise alignment via hyperalignment (error bars, 95% C.I. from 20 individual pairs). It is worth mentioning that optimal transport achieved higher conversion accuracies but lower DNN feature decoding accuracies.

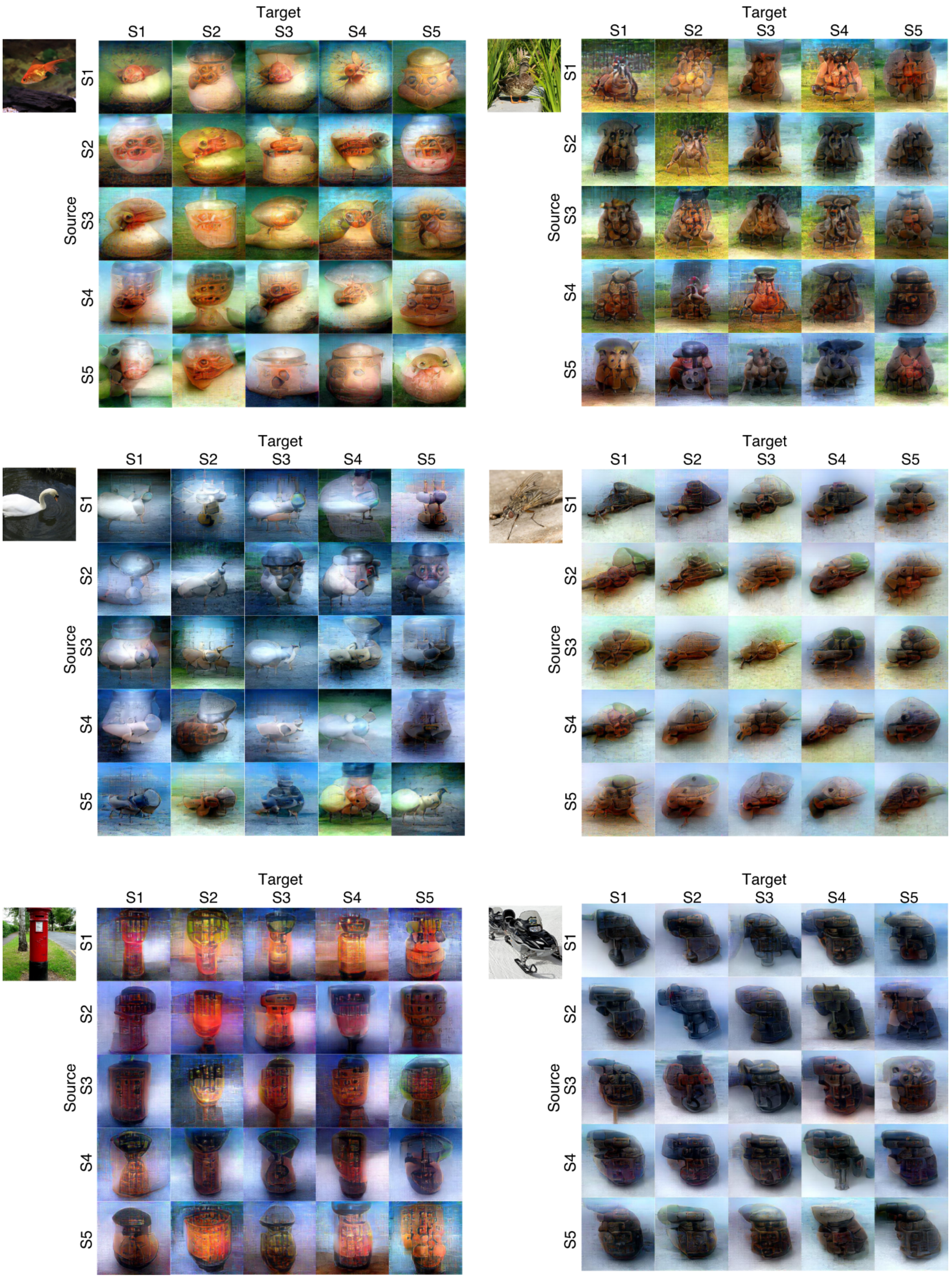
Target
S1 S2 S3 S4 S5
Source
S1
S2
S3
S4
S5

**Fig. S7. Reconstructed natural images across individual pairs.** For each image, the diagonal images in each block are the reconstructed images in the Within condition; the off-diagonal images are the reconstructed images in the Across-functional condition with the converters trained on 2,400 training samples. All images were reconstructed from the whole visual cortex (VC).

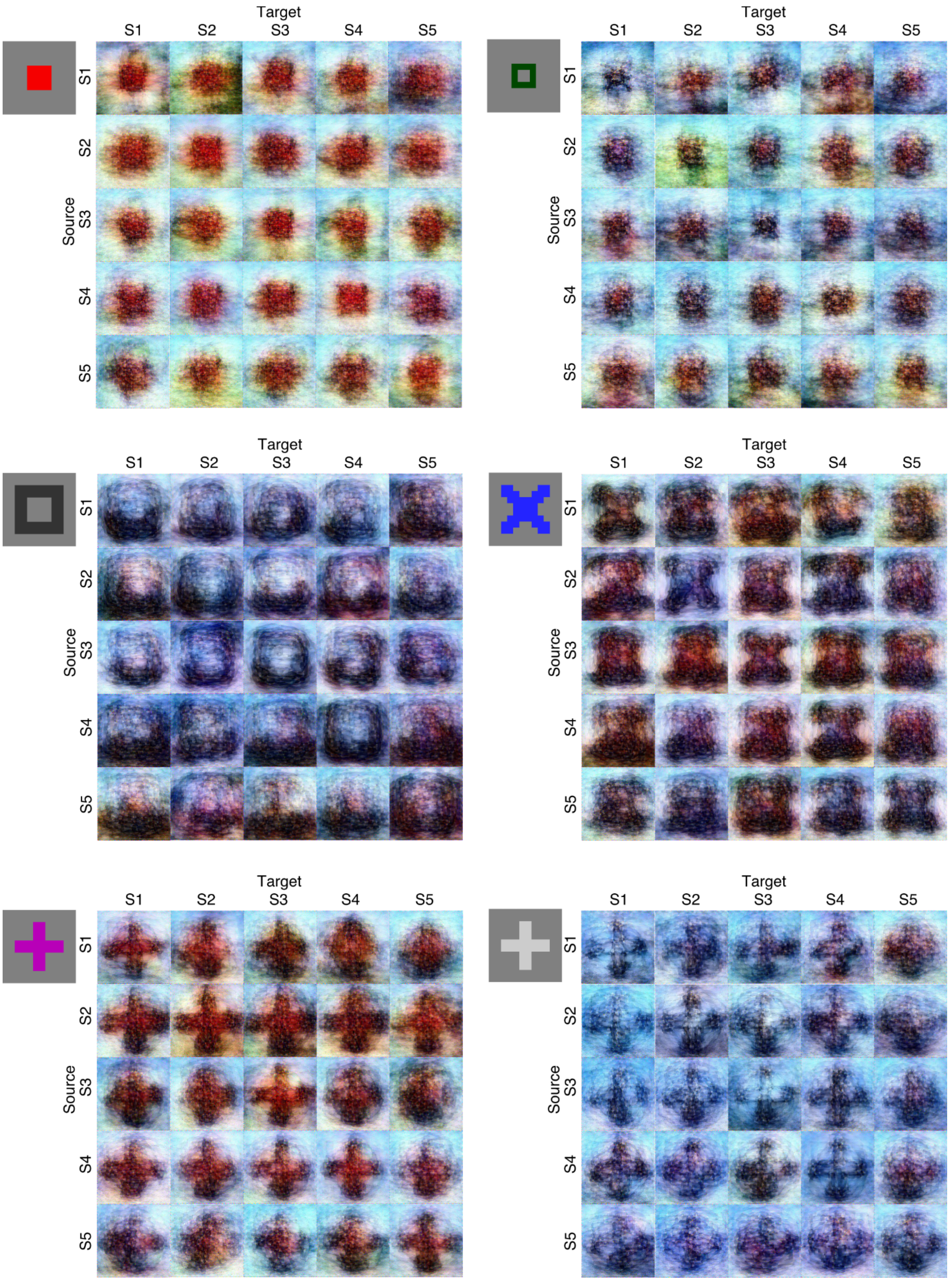
Target
S1
S2
S3
S4
S5
Source
S1
S2
S3
S4
S5

**Fig. S8. Reconstructed artificial images across individual pairs.** For each image, the diagonal images in each block are the reconstructed images in the Within condition, while the off-diagonal images are the reconstructed images in the Across-functional condition with the converters trained on 2,400 training samples. All images were reconstructed from the whole visual cortex (VC).

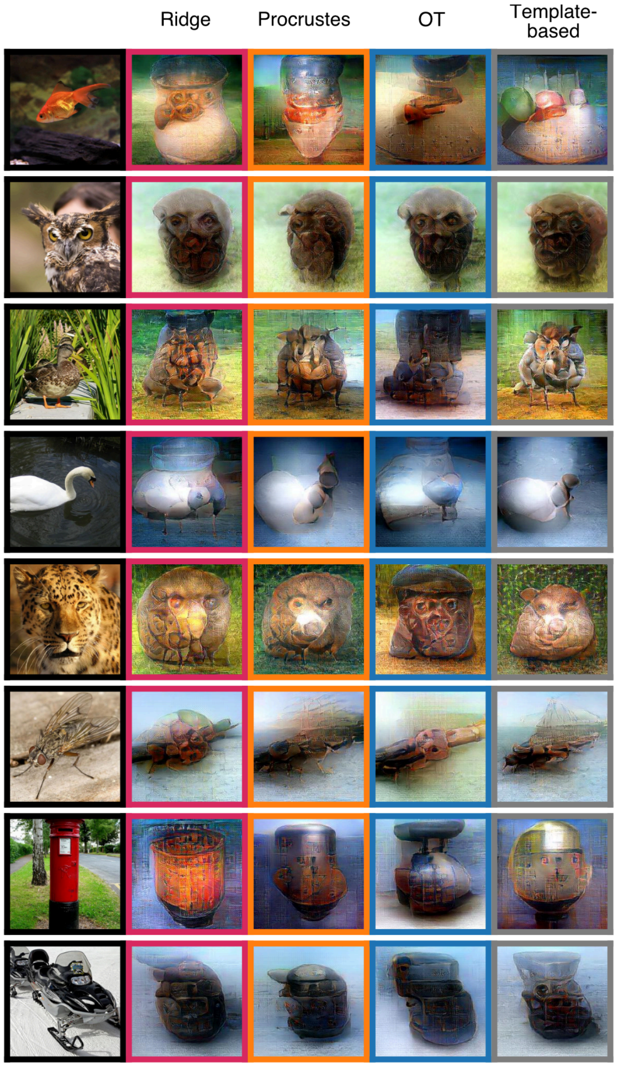


**Fig. S9. Reconstructed images obtained via different methods of functional alignment.** The images were reconstructed with Ridge-based neural code converter, Procrustes transformation, optimal transport (OT), and template-based pairwise alignment via hyperalignment. All images were reconstructed from the whole visual cortex (VC).

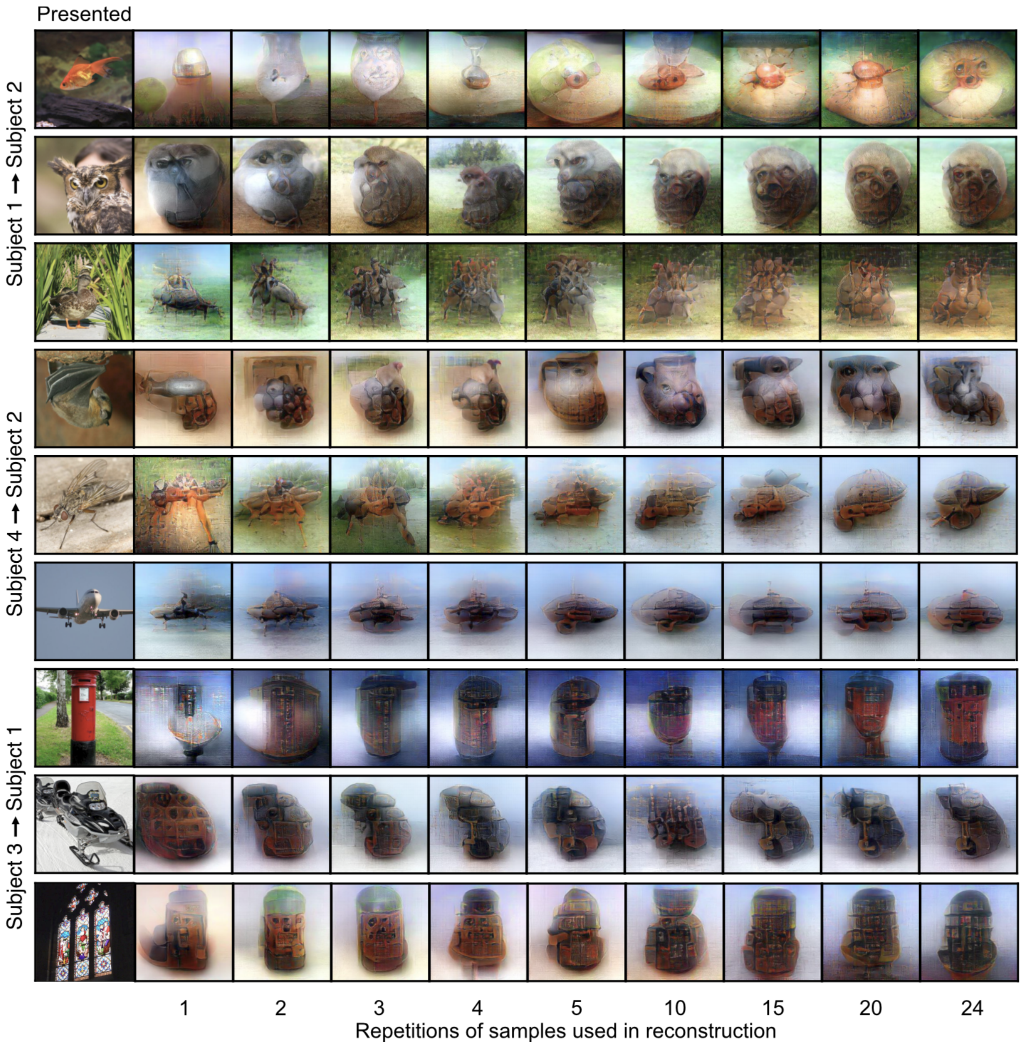


**Fig. S10. Reconstructed images using different repetitions of samples.** The converted fMRI samples corresponding to a visual image were averaged over repetitions and were reconstructed into an image. The reconstructed images were shown for three representative individual pairs. All images were reconstructed from the whole visual cortex (VC).

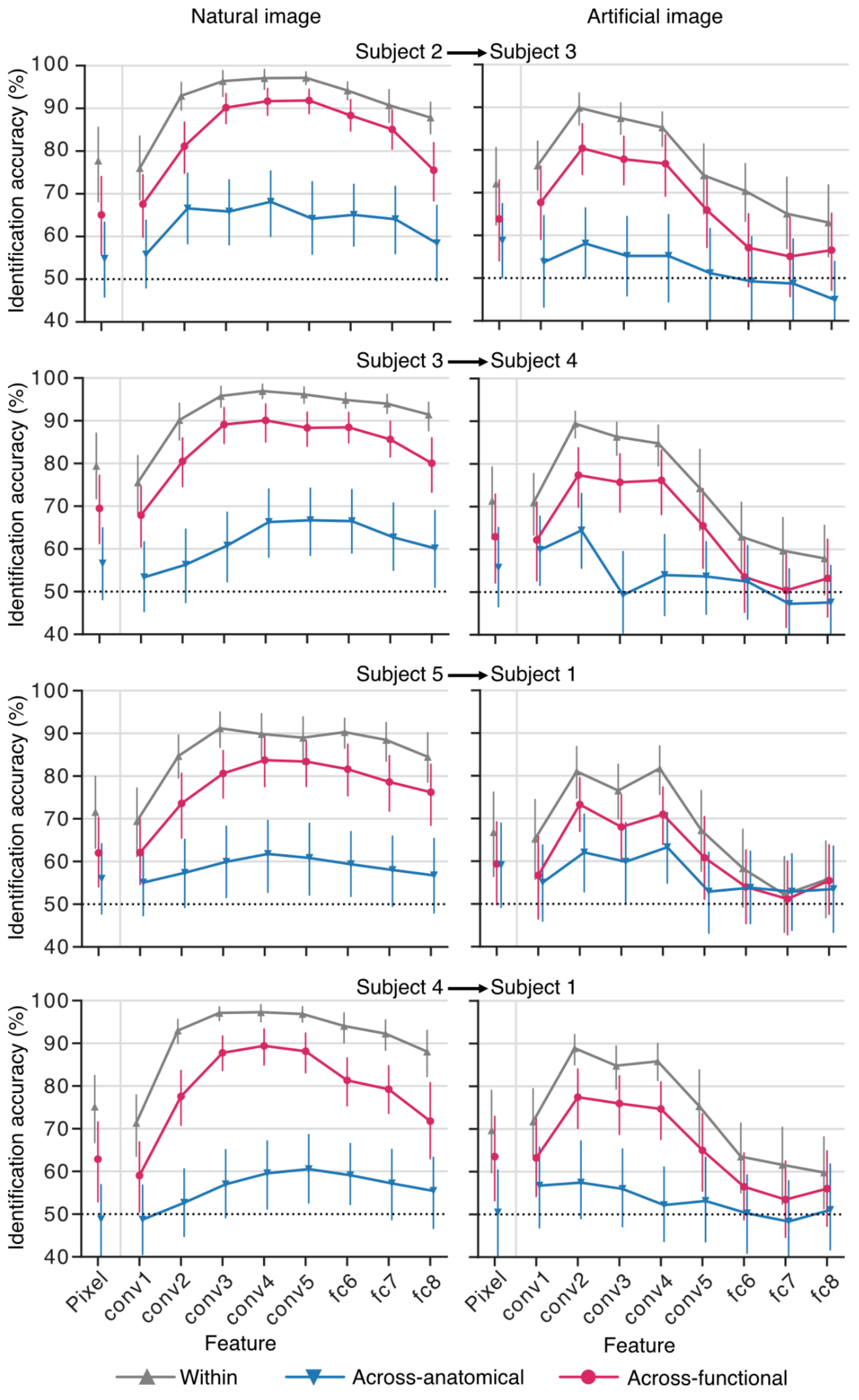


**Fig. S11. Identification accuracies of representative individual pairs.** The identification accuracies for 50 reconstructed natural images were used to calculate a mean identification accuracy and its 95% confidence interval for the Within, Across-anatomical, and Across-functional conditions (left, natural images; right, artificial images; error bar, 95% C.I. of identification accuracies across reconstructed images; dotted lines, chance level = 50%).

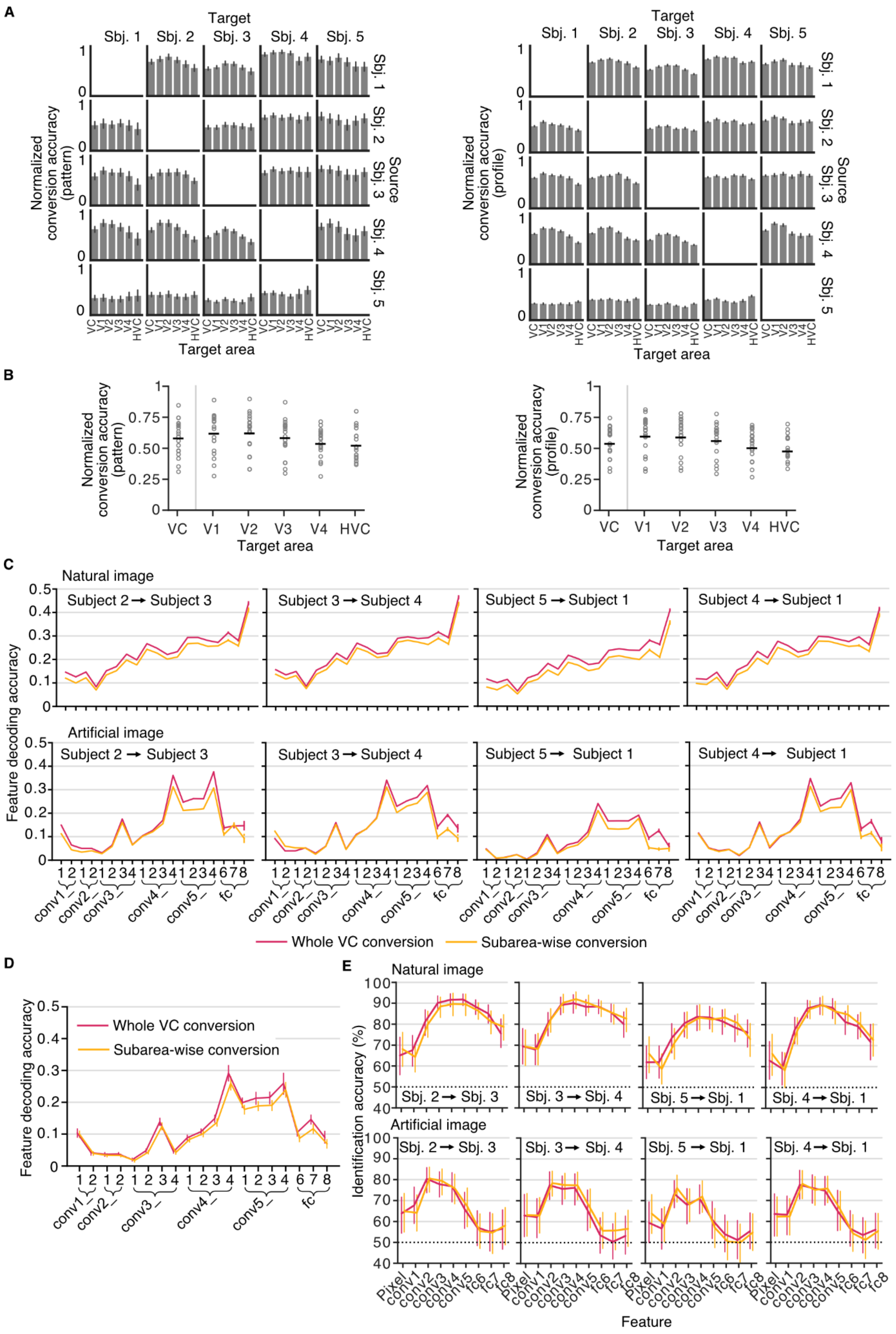
A
Target
Sbj. 1
Sbj. 2
Sbj. 3
Sbj. 4
Sbj. 5
Source
Normalized conversion accuracy (pattern)
Normalized conversion accuracy (profile)
VC V1 V2 V3 V4 HVC
Target area
B
Normalized conversion accuracy (pattern)
Normalized conversion accuracy (profile)
0 0.25 0.50 0.75 1
VC V1 V2 V3 V4 HVC
Target area
C
Natural image
Artificial image
Subject 2 → Subject 3
Subject 3 → Subject 4
Subject 5 → Subject 1
Subject 4 → Subject 1
Feature decoding accuracy
0 0.1 0.2 0.3 0.4 0.5
conv1_ conv2_ conv3_ conv4_ conv5_ fc
Whole VC conversion
Subarea-wise conversion
D
Feature decoding accuracy
Whole VC conversion
Subarea-wise conversion
conv1_ conv2_ conv3_ conv4_ conv5_ fc
E
Natural image
Artificial image
Sbj. 2 → Sbj. 3
Sbj. 3 → Sbj. 4
Sbj. 5 → Sbj. 1
Sbj. 4 → Sbj. 1
Identification accuracy (%)
40 50 60 70 80 90 100
Pixel conv1 conv2 conv3 conv4 conv5 fc6 fc7 fc8
Feature

**Fig. S12. Evaluations of subarea-wise neural code converters.**

(A) Conversion accuracies of individual pairs. The pattern correlation coefficients for 50 visual stimuli were used to calculate a mean conversion accuracy (pattern) and its 95% confidence interval. The profile correlation coefficients for voxels were used to calculate a mean conversion accuracy (profile) and its 95% confidence interval (right; error bars on left panel, 95% C.I. across visual images; error bars on the right panel, 95% C.I. across voxels).

(B) Distributions of normalized pattern or profile correlation coefficients across 20 individual pairs are shown for VC and visual subareas. Each horizontal black dash indicates the mean value over 20 individual pairs; each circle represents the correlation coefficients of an individual pair.

(C) DNN feature decoding accuracy of natural images and artificial images for four representative individual pairs. The decoding accuracies for all DNN units in each layer were used to calculate a mean decoding accuracy and its 95% confidence interval (error bars, 95% C.I. across DNN units).

(D) DNN feature decoding accuracy of artificial images for 20 individual pairs averaged (error bars, 95% C.I. from 20 individual pairs).

(E) Identification accuracies with the reconstructed natural and artificial images for four representative individual pairs. The identification accuracies for individual reconstructed images were used to calculate a mean identification accuracy and its 95% confidence interval. For the natural images, 2/20 pairs showed higher significant accuracies for the subarea-wise conversion; 6/20 pairs showed higher significant accuracies for the whole conversion; for the artificial images, 3/20 pairs showed higher significant accuracies for the subarea-wise conversion; 2/20 pairs showed higher significant accuracies for the whole conversion (ANOVA in individual pairs, see Materials and Methods: “Statistics”).

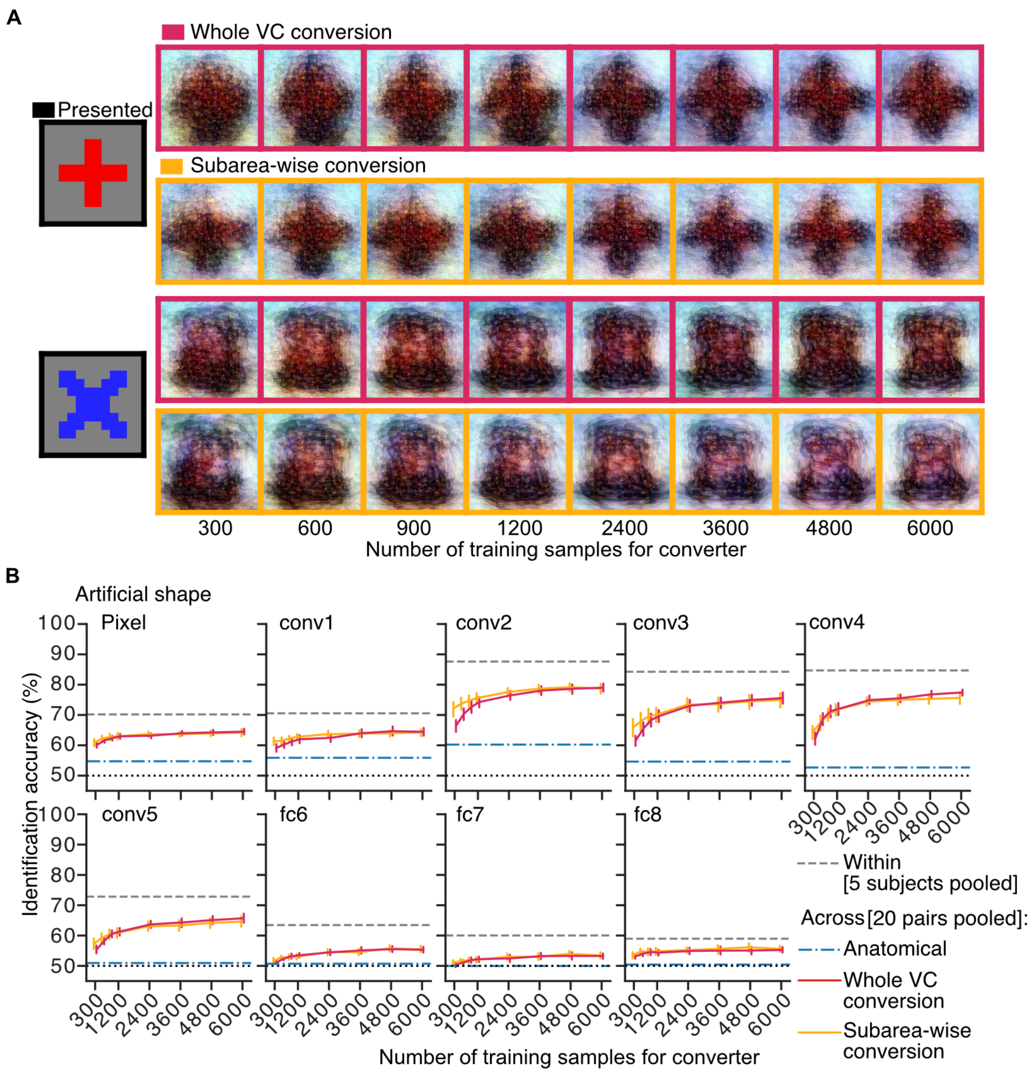


**Fig. S13. Effect of the number of training data for the converter on the reconstruction of artificial images**

(A) Reconstructed images. All reconstructed images were produced from the same subject pair (source, Subject 3; target, Subject 1)

(B) Identification accuracy. Identification accuracies were calculated with the pixel values and the extracted DNN feature values (AlexNet) from the reconstructed artificial images with varying numbers of training data for the whole VC and subarea-wise converters. The results are shown with those from the within-individual condition (Within) and the anatomical alignment (Across-anatomical). The identification accuracies for 20 individual pairs were used to calculate a mean identification accuracy and its 95% confidence interval. The subarea-wise and whole VC conversions showed similar accuracies with more than 900 training samples, but the subarea-wise conversion outperformed the whole VC conversion with 900 or fewer training samples (ANOVA within individual pairs, effect of conversion type, $p < .05$ in 10, 8, 5, 2, 2, 1, 1, and 2 out of 20 pairs for the eight training sample numbers, respectively; group analysis on the mean accuracies of individual pairs, $p < .05$ at 300, 600, and 900 sample; Bonferroni-corrected by eight; error bars, 95% C.I. from 20 individual pairs for whole VC and subarea-wise conversion; dotted lines, chance level = 50%).

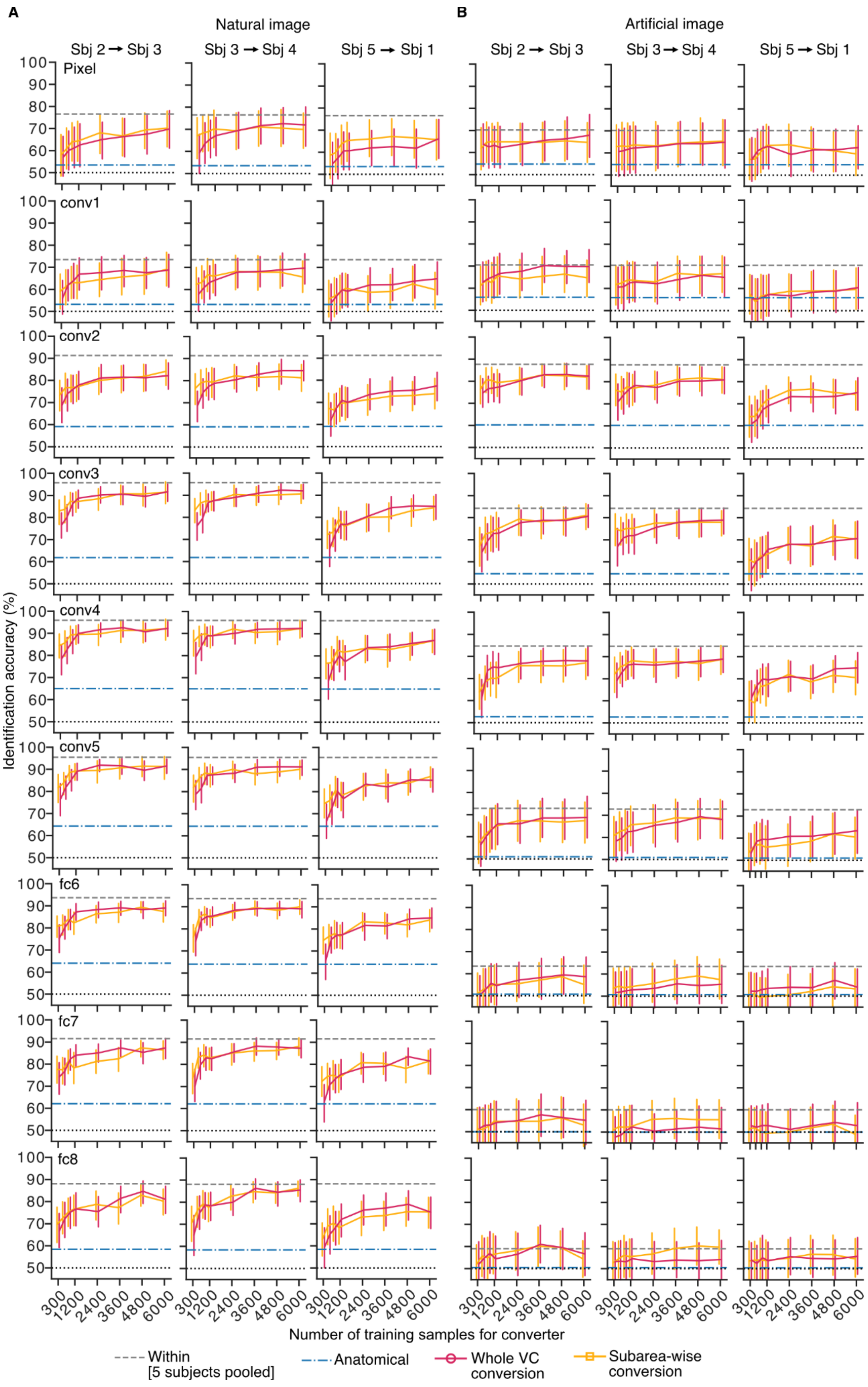
A
B
Natural image
Artificial image
Sbj 2 → Sbj 3
Sbj 3 → Sbj 4
Sbj 5 → Sbj 1
Pixel
conv1
conv2
conv3
conv4
conv5
fc6
fc7
fc8
Identification accuracy (%)
100 90 80 70 60 50
300 1200 2400 3600 4800 6000
Number of training samples for converter
Within [5 subjects pooled]
Anatomical
Whole VC conversion
Subarea-wise conversion

**Fig. S14. Identification accuracies of representative individual pairs with different numbers of training samples for converters.**

(A) Identification accuracies with the reconstructed natural images of three representative individual pairs. For each feature and each training sample condition, the identification accuracies for 50 reconstructed images were used to calculate a mean identification accuracy and its 95% confidence interval. Subarea-wise conversion only out performed whole VC conversion when the training sample is 900 or fewer training samples (ANOVA in individual pairs, effect of conversion type, $p < .05$ in 18, 10, 4, 3 , 2, 1, 4, and 1 out of 20 pairs for the eight training sample numbers; error bar, 95% C.I. across reconstructed images; dotted lines, chance level = 50%).

(B) Identification accuracies with the reconstructed artificial images of three representative individual pairs are shown. For each feature and a training sample condition, the identification accuracies for 40 reconstructed images were used to calculate a mean identification accuracy and its 95% confidence interval. Subarea-wise conversion only out performed whole VC conversion when the training sample is 900 or fewer training samples (ANOVA in individual pairs, effect of conversion type, $p < .05$ in 10, 8, 5, 2, 2, 1, 1, and 2 out of 20 pairs for the eight training sample numbers; error bar, 95% C.I. across reconstructed images; dotted lines, chance level = 50%).

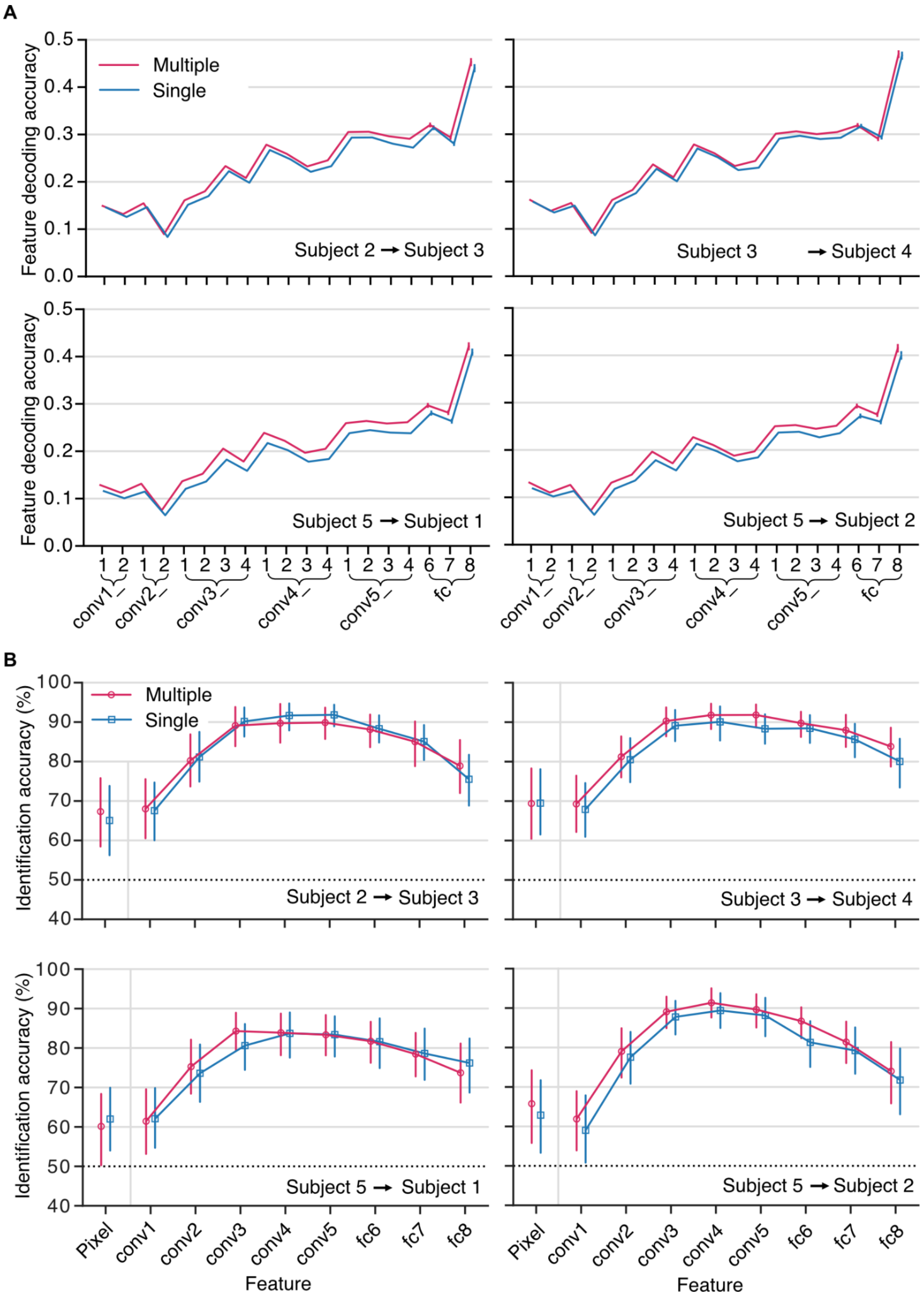


**Fig. S15. Feature decoding and identification analyses of representative individual pairs with multiple- and single-subject feature decoders.**

(A) DNN feature decoding accuracies of natural images of four representative individual pairs. The decoding accuracies for all DNN units in each layer were used to calculate a mean decoding accuracy and its 95% confidence interval (error bars; 95% C.I. across DNN units).

(B) Identification accuracies with the reconstructed natural images of four representative individual pairs. The identification accuracies for 50 reconstructed natural images were used to calculate a mean identification accuracy and its 95% confidence interval. The multiple-subject feature decoders outperformed the single-subject feature decoders (ANOVA, effect of decoder type with DNN layers as between-subject factor, $p < .05$ in 10/20 individual pairs; error bars, 95% C.I. across reconstructed images; dotted line, chance level = 50%).

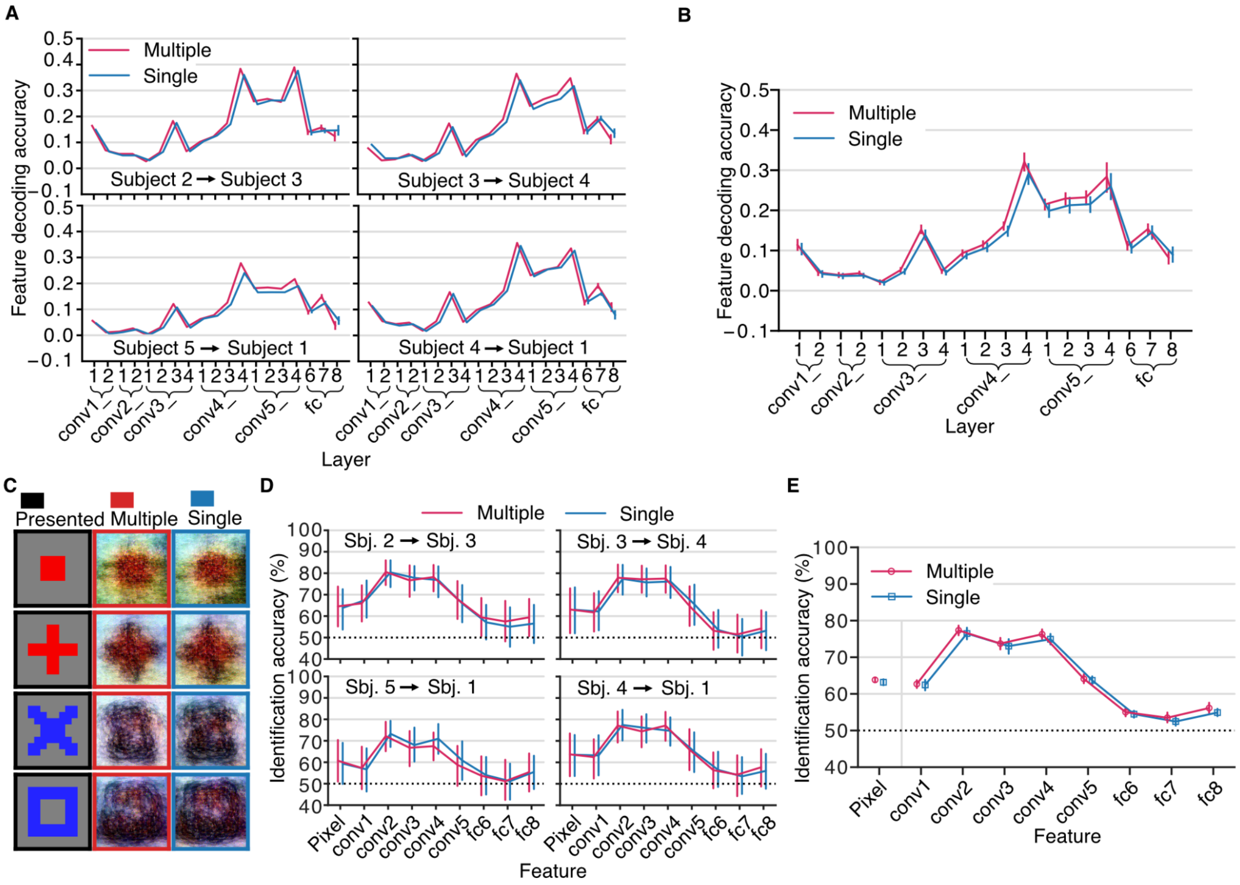


**Fig. S16. Evaluation of multiple subject feature decoders using artificial images dataset.**

(A) DNN feature decoding accuracies of four representative individual pairs for artificial images. The multiple-subject feature decoders were trained on pooled data (24,000 samples); the single-subject feature decoders were trained on only one single subject's data (6,000 samples). The converters were trained with 2,400 samples. The decoding accuracies for all DNN units in each layer were used to calculate a mean decoding accuracy and its 95% confidence interval (error bars, 95% C.I. across DNN units).

(B) DNN feature decoding accuracy averaged over 20 individual pairs. The decoding accuracies for 20 individual pairs in each layer were used to calculate a mean decoding accuracy and its 95% confidence interval. The multiple-subject feature decoders outperformed the single-subject feature decoders (ANOVA, effect of decoder type, $F(1, 361) = 172$, $p < .001$, $\eta_p^2 = .32$; error bars, 95% C.I. from 20 individual pairs).

(C) Reconstructed artificial images for the multiple- and single-subject conditions.

(D) Identification accuracies with the reconstructed artificial images for four representative individual pairs. The identification accuracies for 40 reconstructed artificial images were used to calculate a mean identification accuracy and its 95% confidence interval. The multiple-subject feature decoders outperformed the single-subject feature decoders in 6 out of 20 individual pairs (ANOVA, effect of decoder type, $p < .05$; error bars, 95% C.I. across reconstructed images).

(E) Identification accuracy averaged over 20 individual pairs. The identification accuracies for 20 individual pairs were used to calculate a mean identification accuracy and its 95% confidence interval. The multiple-subject feature decoders outperformed the single-subject feature decoders at the group level (ANOVA; effect of decoder type on group level, $F(1, 171) = 35.9$, $p < .001$, $\eta_p^2 = .17$; error bars, 95% C.I. from 20 individual pairs; dotted lines, chance level = 50%).

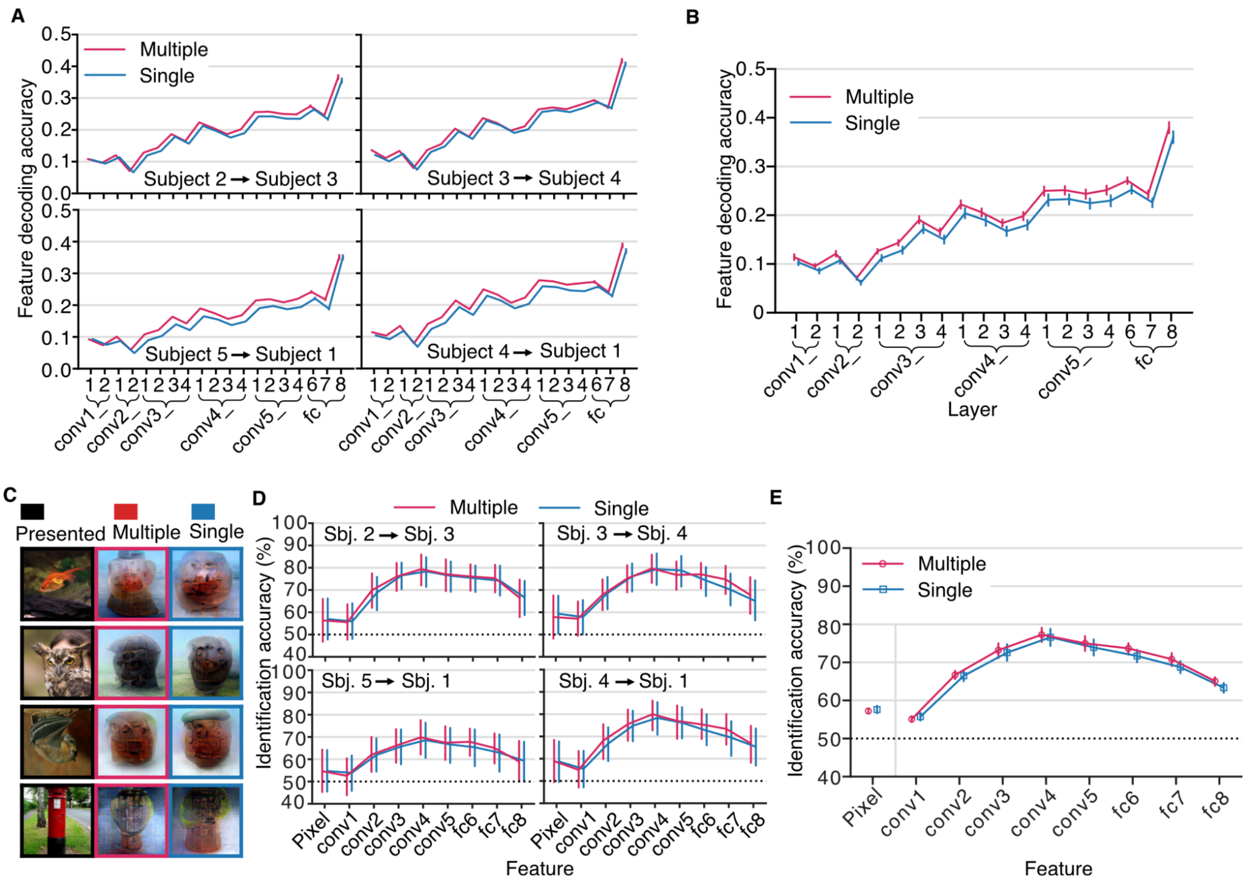


**Fig. S17. Pooling data analysis with limited data for a novel subject.**

(A) DNN feature decoding accuracies of four representative individual pairs for natural images (*c.f.* Fig. S13). The converters were trained with 300 samples. The decoding accuracies for all DNN units in each layer were used to calculate a mean decoding accuracy and its 95% confidence interval (error bars, 95% C.I. across DNN units).

(B) DNN feature decoding accuracy averaged over 20 individual pairs. The decoding accuracies for 20 individual pairs in each layer were used to calculate a mean decoding accuracy and its 95% confidence interval. The multiple-subject feature decoders outperformed the single-subject feature decoders (ANOVA, effect of decoder type, $F(1, 361) = 1300$, $p < .001$, $\eta_p^2 = .78$; error bars, 95% C.I. from 20 subject pairs).

(C) Reconstructed natural images for the multiple- and single-subject conditions.

(D) Identification accuracies of the reconstructed natural images for four representative individual pairs. The identification accuracies for 50 reconstructed natural images were used to calculate a mean identification accuracy and its 95% confidence interval. The multiple-subject feature decoders outperformed the single-subject feature decoders in 5 out of 20 individual pairs

(ANOVA, effect of decoder type, $p < .05$; error bars, 95% C.I. across reconstructed images).
(E) Identification accuracies averaged over 20 individual pairs. The identification accuracies for 20 individual pairs were used to calculate a mean identification accuracy and its 95% confidence interval. The multiple-subject feature decoders outperformed the single-subject feature decoders at the group level (ANOVA; effect of decoder type, $F(1, 171) = 28.3$, $p < .001$, $\eta_p^2 = .14$; error bar, 95% C.I. from 20 subject pairs; dotted lines, chance level = 50%).

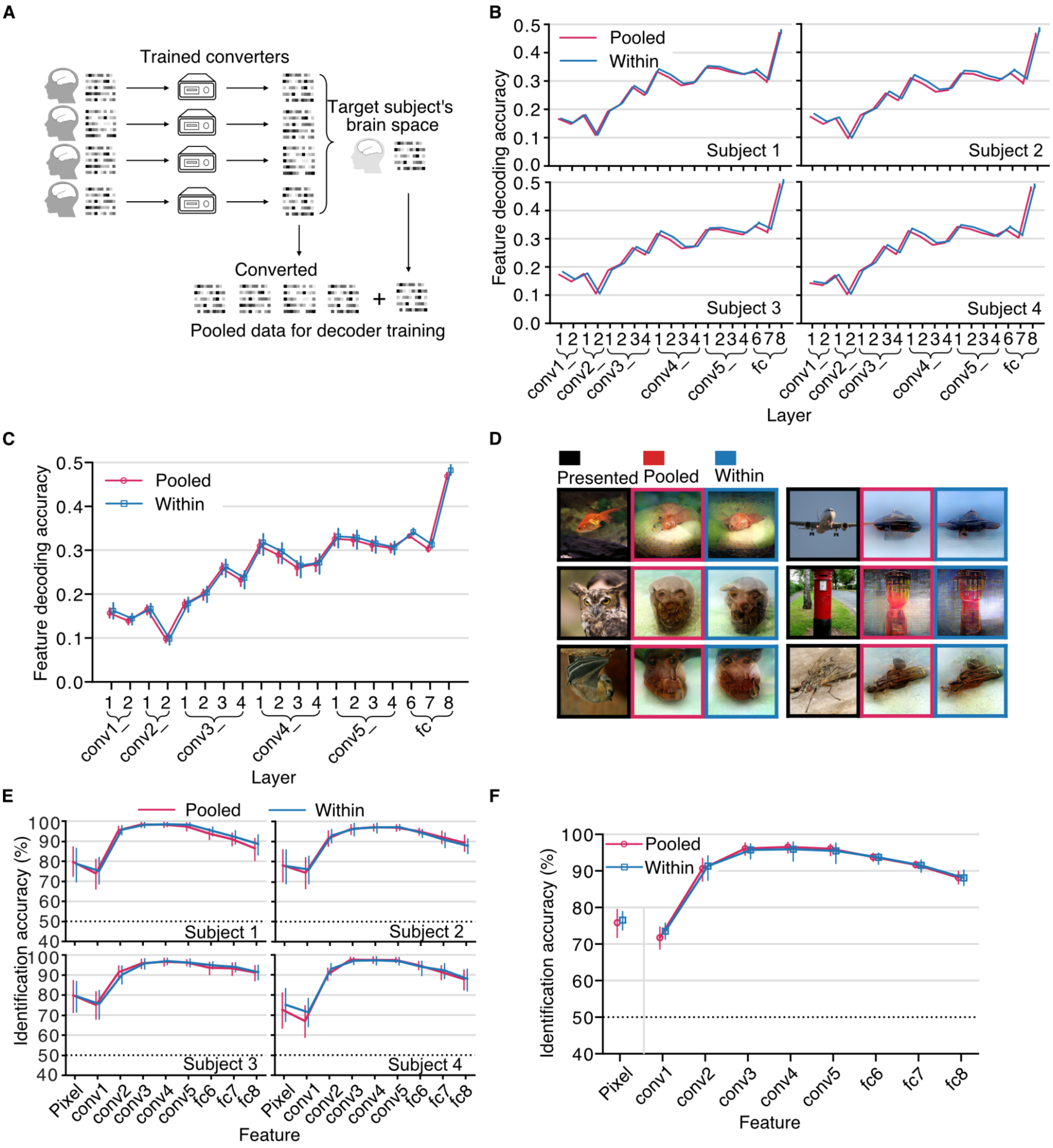


**Fig. S18. Augmenting data of a subject.**

(A) Illustration of the augmenting data procedure. This analysis examined the effect of augmenting a subject's data by pooling other subjects' data into a subject's brain space on the performance of the feature decoders and visual image reconstruction. One subject was selected as the target subject and the data (6,000 samples per subject) of the other subjects were pooled into the target brain space via the converters trained with 2,400 samples. We then trained the

DNN feature decoders with the pooled data (5 × 6,000 = 30,000 samples). The decoders are called “pooled data feature decoders.” We then tested the “pooled data feature decoders” using the target subject’s data to compare with the “within-individual feature decoders,” which were trained with 6,000 samples of individual data. The decoders were evaluated by DNN feature decoding and visual image reconstruction. The same analysis was repeated by iterating the target subject among the five subjects.

(B) DNN feature decoding accuracies of four representative subjects for natural images. The decoding accuracies for all DNN units in each layer were used to calculate a mean decoding accuracy and its 95% confidence interval (error bars, 95% C.I. across DNN units).

(C) DNN feature decoding accuracy averaged over five subjects. The decoding accuracies for five subjects were used to calculate a mean decoding accuracy and its 95% confidence interval. The pooled data feature decoders slightly underperformed the within-individual feature decoders (ANOVA, effect of decoder type, $F(1, 76) = 207$, $p < .001$, $\eta_p^2 = .73$; error bars, 95% C.I. from five subjects).

(D) Reconstructed natural images from the pooled data feature decoders and the within-individual feature decoders.

(E) Identification accuracies with the reconstructed natural images for four representative subjects. The identification accuracies for individual reconstructed images were used to calculate a mean identification accuracy and its 95% confidence interval. There were no statistical differences in identification accuracies between two conditions for Subject 2, 3, and 5 while the pooled data feature decoders slightly outperformed within-individual feature decoders for Subject 1 and 4 (ANOVA, effect of decoder type, $p < .05$; error bars, 95% C.I. across reconstructed images).

(F) Identification accuracies with the reconstructed natural images averaged over five subjects. The identification accuracies for five subjects were used to calculate a mean identification accuracy and its 95% confidence interval. There was no statistical difference in identification accuracies between the pooled data feature decoders and the within-individual feature decoders at the group level (ANOVA, $F(1, 36) = .40$, $p = .53$, $\eta_p^2 = .011$; error bar, 95% C.I. from five subjects; dotted lines, chance level = 50%).

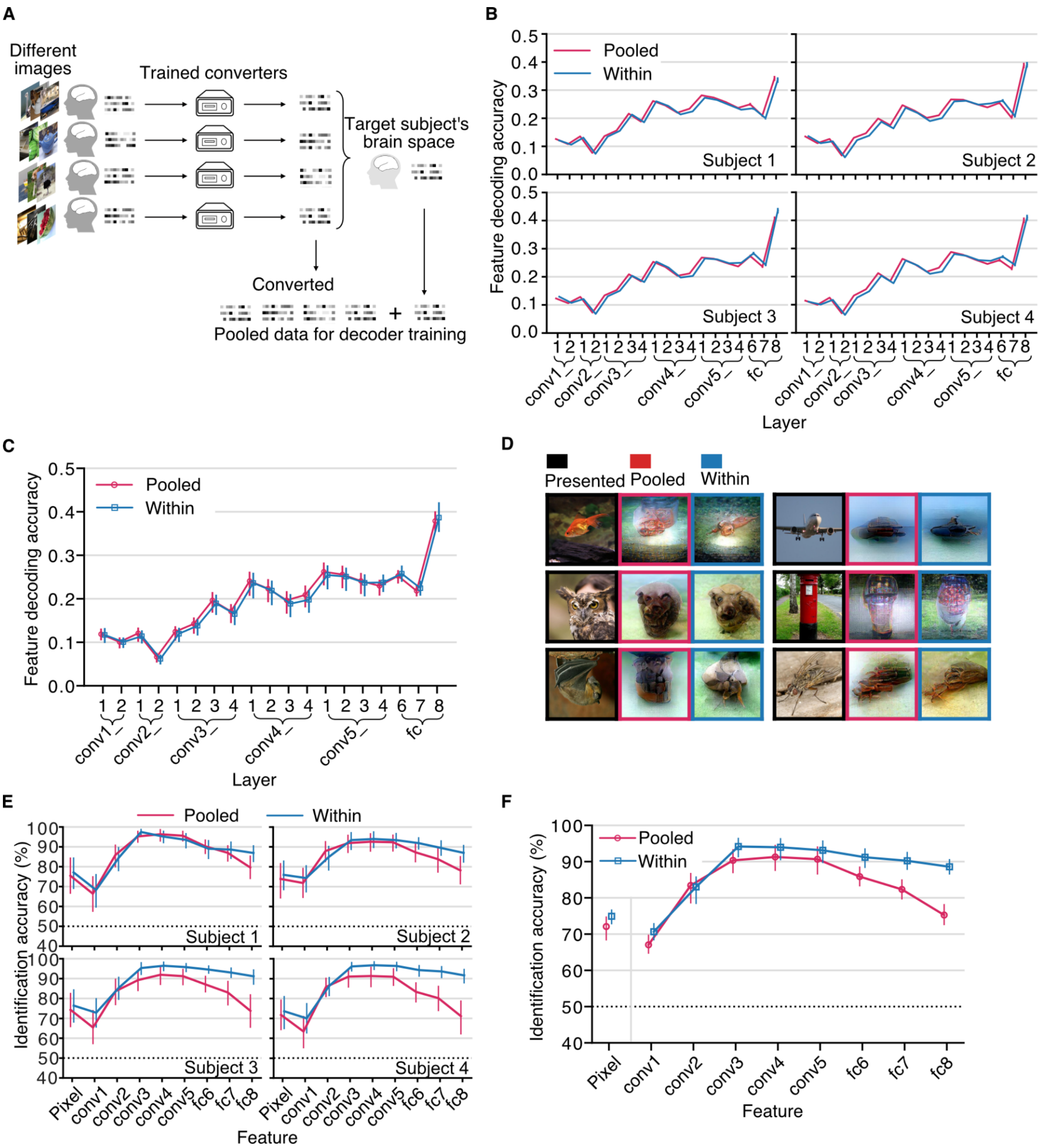


**Fig. S19. Augmenting data of a subject with stimulus variation.**

(A) Illustration of the pooling data procedure. This analysis examined the effect of increasing the stimulus variability by pooling data of other subjects viewing different sets of images into a subject's brain space on the performance of the feature decoders and visual image reconstruction. The 1,200 natural images were split into six sets of 200 images: the first set is the common image set; each subject has the 1,000 fMRI data samples (200 images × 5 repetitions) of the common images; additionally, each subject was assigned one of the remaining five image sets, and was assumed to have the 1,000 fMRI data samples (200 images × 5 repetitions) of the image set. We trained the within-individual DNN feature decoders on these 2,000 samples. Then, each subject was iteratively selected as the target subject and the data of uncommon image sets of the other subjects were pooled into the target brain space via the converters trained with the 1,000 samples of the common image set. We trained the DNN feature decoders with the pooled 6,000 samples and called them the "pooled data feature decoders." The decoders were evaluated by DNN feature decoding and visual image reconstruction.

(B) DNN feature decoding accuracies of four representative subjects for natural images. The decoding accuracies for all DNN units in each layer were used to calculate a mean decoding accuracy and its 95% confidence interval (error bars, 95% C.I. across DNN units).

(C) DNN feature decoding accuracies of natural images averaged over five subjects. The decoding accuracies for five subjects were used to calculate a mean decoding accuracy and its 95% confidence interval. The pooled data feature decoders marginally outperformed the within-individual feature decoders (ANOVA, effect of decoder type, $F(1, 76) = 22.3$, $p < .001$, $\eta_p^2 = .23$; error bars, 95% C.I. from five subjects).

(D) Reconstructed natural images from the pooled data feature decoders and the within-individual feature decoders.

(E) Identification accuracies with the reconstructed natural images for four representative subjects. The identification accuracies for 50 reconstructed natural images were used to calculate a mean identification accuracy and its 95% confidence interval. The pooled data feature decoders underperformed the within individual feature decoders, with four subjects showing statistical significances (ANOVA, effect of decoder type, $p < .05$; error bars, 95% C.I. across reconstructed images).

(F) Identification accuracies with the reconstructed natural images averaged over five subjects. The identification accuracies for five subjects were used to calculate a mean identification accuracy and its 95% confidence interval. The pooled data feature decoders underperformed the within individual feature decoders at the group level (ANOVA, effect of decoder type, $F(1, 36) = 78.7$, $p <.001$, $\eta_p^2 = .69$). The results suggested that the current converter models could not fully resolve the feature mismatch of brain activity patterns between individuals, and the decoder struggled to learn the statistical relationships between DNN features and brain activity patterns with the existence of multiple subjects (error bar, 95% C.I. from five subjects; dotted lines, chance level = 50%).

**Video S1. Deep image reconstruction in within-individual and across-individual conditions.** The iterative optimization process is shown (top left, presented images; top right, Within condition; bottom left, Across-anatomical condition; bottom right, Across-functional condition).

**Video S2. Deep image reconstruction via converters trained with varying numbers of data.** The iterative optimization process is shown (top left, presented images; top right, 320 mins of data; bottom left, 160 mins of data; bottom right, 40 mins of data).
